# Using Extracellular Vesicles Released by GDNF-transfected Macrophages for Therapy of Parkinson’s Disease

**DOI:** 10.1101/2022.05.25.493424

**Authors:** Yuling Zhao, Matthew J. Haney, John K. Fallon, Myosotys Rodriguez, Carson J. Swain, Camryn J. Arzt, Philip C. Smith, Matthew Shane Loop, Emily B. Harrison, Nazira El-Hage, Elena V. Batrakova

## Abstract

Extracellular vesicles (EVs) are cell-derived nanoparticles that facilitate transport of proteins, lipids and genetic material playing important roles in intracellular communication. They have a remarkable potential as non-toxic and non-immunogenic nanocarriers for drug delivery to unreachable organs and tissues, in particular, the central nervous system (CNS). Herein, we developed a novel platform based on macrophage derived EVs to treat Parkinson’s disease (PD). Specifically, we evaluated the therapeutic potential of EVs secreted by autologous macrophages that were transfected *ex vivo* to express glial cell line-derived neurotrophic factor (GDNF). EV-GDNF were collected from conditioned media of GDNF-transfected macrophages and characterized for GDNF content, size, charge, and expression of EV-specific proteins. The data revealed that along with the encoded neurotrophic factor, EVs released by pre-transfected macrophages carry GDNF-encoding DNA. Four months-old transgenic Parkin Q311(X)A mice were treated with EV-GDNF *via* intranasal administration, and the effect of this therapeutic intervention on locomotor functions was assessed over a year. Significant improvements in mobility, increase in neuronal survival, and decrease in neuroinflammation were found in PD mice treated with EV-GDNF. No offsite toxicity caused by EV-GDNF administrations was detected. Overall, EV-based approach can provide a versatile and potent therapeutic intervention for PD.

## 1. Introduction

The National Parkinson Foundation^®^ estimates that Parkinson’s disease (PD) affects more than a million individuals in the U.S. with up to 60,000 new cases diagnosed each year. One of the greatest challenges is to provide efficient treatment for PD besides the replacement of neurotransmitters. Loss of dopaminergic neurons in the *Substantia Nigra pars compacta* (*SNpc)* along with inflammation in the brain and production of the excessive amount of reactive oxygen species (ROS) are the hallmarks of PD. While many potent therapeutic proteins, including antioxidants and neurotrophic factors were identified, the blood brain barrier (BBB) remains a seemingly insurmountable obstacle to the routine use of systemically administered macromolecules. Contrary to stroke [1], diabetes [2], or traumatic brain injury [3] that can severely disrupt the BBB, PD patients usually do not have substantial changes in the BBB permeability. Nevertheless, later advances indicate that some BBB disfunction may be developed upon late stages of neurodegenerative disorders, including PD dementia [2].

Different strategies are currently being developed to improve drug delivery across the BBB. They can be divided into two principal groups: the increasing drug influx into the brain and restricting drug efflux out of the brain. Approaches for the first group include: (*i*) modification of a drug chemical structure, for example, increasing lipophilicity of the molecule [4]; or (*ii*) using carrier-mediated and receptor-mediated transcytosis [5]; or (*iii*) incorporation of a drug into micro- and nano-containers, including EVs [6]. The second group that is focused on the restricting drug efflux includes co-administration of competitive or non-competitive inhibitors of drug efflux transporters such as P-glycoprotein (Pgp), multidrug resistance protein (MRP), breast cancer resistance protein (BCRP), and the multi-specific organic anion transporter (MOAT), the members of the ABC cassette (ATP-binding cassette) family [5, 7].

In this work, we utilized strategy that belongs to the first group using EVs for transport of a potent neurotrophic factor, GDNF, and provided prove-of-concept in a transgenic mouse PD model, Parkin-Q311X(A) mice. GDNF is known to promote neuronal survival that is essential to mitigate neurodegeneration during PD [8-13]. Furthermore, a restoration and regeneration of dopaminergic neurons by activated macrophages and microglia that overexpress GDNF was demonstrated earlier during natural healing process in injured striatum [14]. Regrettably, current attempts to deliver GDNF to the CNS have been hampered by pharmacological issues including poor penetration across the BBB and offsite toxicity upon systemic administration. In recent clinical investigations, direct infusions of GDNF to the affected brain areas were shown to produce promising therapeutic effects in PD patients [15]. However, these brain infusions carry a high risk of adverse effects and have poor patient adherence. As such, the development of innovative strategies that allow efficient GDNF delivery to the brain is of great importance.

In this regard, the field of nanotechnology holds enormous promise for the development of versatile drug delivery systems. The incorporation of drugs into nanocarriers would protect them against degradation and elimination in the bloodstream, and target therapeutics to a disease site. Much effort has been dedicated to the development of nanoformulations for drug delivery [16, 17], but these efforts have been met with limited success. In fact, the opsonization of drug-loaded synthetic nanoparticles in the bloodstream caused two main problems, nanotoxicity and rapid drug clearance by mononuclear phagocyte system (MPS). To circumvent this problem, we engineered EV-based biomimetic delivery system capable of targeted transport of this neurotrophic factor to the brain. Using EVs as beneficial bio-nanostructures has a potential to overcome the major drawbacks related to opsonization and cytotoxicity and introduce them for clinical applications. Composed of cellular membranes with multiple adhesive proteins on their surface, EVs are known to specialize in cell-cell communication facilitating transport of their cargo to target cells. It was reported that EVs penetrate the BBB and can deliver incorporated therapeutics upon systemic and loacal administration [18-21]. Three different mechanisms of EVs transport across the BBB were suggested: receptor-mediated transcytosis, lipid-raft mediated endocytosis, and macropinocytosis [20]. In addition, EVs exert unique biological activity reflective of their origin that allow to capitalize on specific properties of parent cells and amplify therapeutic effects of EV-based formulations [22]. These exceptional features of EVs should work in concert to dramatically improve the therapeutic efficacy of current treatment strategies utilizing GDNF.

In our earlier investigations, we assessed several types of parent cells and selected inflammatory response cells, macrophages that allow targeted drug delivery to the inflamed brain in PD animals. Of note, using macrophages as parent cells is crucial for targeting EVs to inflamed brain tissues. We reported recently that EVs released by macrophages can accumulate in inflamed brain tissues in PD mice at greater amounts than those released by neurons or astrocytes [23]. Specifically, we demonstrated a remarkable ability of immunocyte derived EVs to interact with recipient cells [24-26], target inflamed brain tissues *via* LFA1/ICAM1 interactions, and deliver their therapeutic cargo [22, 27, 28] resulting in profound therapeutic effects in mice with acute neuro-inflammation induced by lipopolysaccharide (LPS), 6-hydroxydopamine (6-OHDA), or 1-methyl-4-phenyl tetrahydropyridine (MPTP). However, these toxin-induced PD models resemble PD at late stages, whereas transgenic animal models are more appropriate to represent early stages of the disease. In this work, we used Parkin-Q311(X)A mice, a transgenic model of PD with a mutation that produces C-terminally truncated parkin. These mice slowly develop degeneration of dopaminergic neurons and progressive decrease in motor activity levels over several months [29].

EVs have already been recognized as promising drug nanocarriers for clinical use. Currently, over 20 clinical trials involving EVs could be fount www.clinicaltrials.gov. Most of them, especially using Mesenchymal Stem Cells (MSCs) derived EVs showed no safety concerns, high feasibility, and significant enhancement of antitumor immune response [30-34]. However, there are some drawbacks, including the upscaling processes of isolation and purification, as well as efficient loading of these natural nanovesicles with various therapeutics. Concerning the manufacture of large lots of EVs nanocarriers, we reported earlier that EVs formulations can be lyophilized and then restored without altering their morphology [24, 25]. More information could be found in the recent review [28].

Regarding methods of drug incorporation into EVs nanocarriers, we developed earlier exogenous loading naïve EVs isolated from parent cells conditioned media. Different techniques were used for loading therapeutics into EVs, including co-incubation, freeze-thaw cycles, sonication, electroporation, extrusion, and permeabilization of EV membranes with saponin [19, 21, 25, 26, 35-37]. As a result, EV-based formulations with a high loading efficacy, sustained drug release, and preservation against degradation and clearance were manufactured. In the present work, we utilized another approach, the endogenous loading EVs with GDNF that was accomplished through the genetic modification of parent cells with GDNF-encoding plasmid DNA (*p*DNA). We characterized the ontained EV-GDNF by ELISA for the levels of GDNF, and RT-PCR for genetic content, as well as by Nanoparticle Tracking Analysis (NTA) and Atomic Force Microscopy (AFM) for size distribution, charge, concentration, and morphology. Furthermore, the presence of specific proteins constitutively expressed in EVs was assessed by Western blot and label free targeted quantitative proteomics. Next, we demonstrated earlier that intranasal (*i*.*n*.) administration route provided high accumulation levels of these nanocarriers in the brain [19]. Therefore, we used this route for EV-GDNF treatment of Parkin Q311(X)A mice.

The novelty of this study can be outlined in three main points. First, we demonstrated that EVs can be loaded with GDNF protein and GDNF-encoding DNA through genetic modification of parent cells, macrophages. Second, the label free targeted quantitative proteomic analysis was applied first time to examine effect of genetic modification of parent cells on the expression and quantification of different integrins that play crucial role on EVs accumulation in target cells. Third, prolonged sustained therapeutic effects followed by *i*.*n*. administrations of EV-GDNF were demonstrated in PD mice over a year. Importantly, multiple lines of evidence for therapeutic efficacy of EV-based formulations were observed, including decreased brain inflammation, significant neuroprotection of dopaminergic neurons in *SNpc* brain region, and improved locomotor functions. Finally, no overall toxicity was detected following administration of macrophage derived EVs.

## 2. Materials and Methods

### 2.1. Plasmids and Reagents

Human GDNF cDNA (NM_199234) was provided by OriGene (Rockville, MD, USA) that was propagated in DH5α E.coli, followed by purification Giga-prep kits (Qiagen, Valencia, CA, USA). The sequence of hGDNF cDNA can be found on web page http://www.origene.com/human_cdna/NM_199234/SC307906/GDNF.aspx, and the plasmid map for GDNF production is presented on **Supplementary Figure S1**. The structure of obtained plasmid was confirmed by electrophoresis with a single cut (AlwNI), and a double cut (AlwNI & PflMI) [38]. Murine macrophage colony-stimulating factor (MCSF) was purchased from Peprotech Inc. (Rocky Hill, NJ, USA). Purified water was obtained from a Picopure® 2 system (Hydro Service and Supplies, Inc. Durham, NC, USA). Trypsin Gold mass spectrometry grade (item # V5280) was purchased from Promega (Madison, WI, USA). Cell culture medium and fetal bovine serum (FBS) were purchased from Gibco Life Technologies, (Grand Island, NY, USA). Solid phase extraction (SPE) cartridges for sample clean up (Strata™-X 33u Polymeric Reversed Phase, 10 mg/mL, part no. 8BS100AAK) were purchased from Phenomenex (Torrance, CA, USA). Fluorescent dyes, 1,1’-Dioctadecyl-3,3,3’,3’-Tetramethylindodicarbocyanine, 4-Chlorobenzenesulfonate Salt (DID) and 4′,6-diamidino-2-phenylindole (DAPI) were purchased from Invitrogen (Carlsbad, CA, USA). Ammonium bicarbonate, dithiothreitol, β-casein (from bovine milk), sodium deoxycholate, iodoacetamide and acetic, formic and trifluoroacetic acids were purchased from Millipore Sigma (St. Louis, MO, USA). Acetonitrile (HPLC grade), methanol and 0.2 mL flat cap PCR tubes (catalog # 14230227) were purchased from Fisher Scientific (Pittsburg, PA, USA). All other chemicals were reagent grade.

### 2.2. Cells

To obtain autologous parental cells for manufacture of GDNF-carrying EVs, female wild type littermates of the transgenic mice (2 mo. old) were used as donors for bone marrow-derived cells that were cultured in 75T flasks over 10 days in the presence of macrophage colony-stimulating factor (MCSF, 1000 U/mL, at 37 ^0^C and 5% CO_2_) [39]. The cells that did not attach to the flask during the culturing were considered as non-differentiated cells and removed with media replaced. The produced primary bone-marrow macrophages (BMM) were characterized by flow cytometry with FACSCalibur (BD Biosciences, San Jose, CA). The obtained data indicated that about 95% of the cells were in fact CD11b+.

### 2.3. Transfection Macrophages

Macrophages were transfected by electroporation. Briefly, 5×107 cells were spanned down at 125 RCF for 5 min, and then re-suspended in 700 µL electroporation buffer (Neon Transfection system, Thermo Fisher Scientific) and supplemented with 30 µg GDNF-*p*DNA. The aliquot of cell suspension with GDNF-*p*DNA (100 µL) was placed into electroporation cell, and electroporated at the following electroporation conditions, outlined in **Supplementary Table S1**. Then, the cells were supplemented with 500 µL antibiotic-free media and cultured for up to 6 days in RPMI 1640 media (Sigma-Aldrich). At each time point, EVs were isolated from the conditioned media, and the GDNF levels in the cells and EVs media were assessed by ELISA as described earlier [40]. Sham macrophages were transfected with sham vector, green fluorescence protein (GFP)-encoding *p*DNA.

### 2.4. EVs Isolation

Concomitant media from GDNF-transfected macrophages was collected, and EVs were isolated using differential centrifugation [41]. First, the cells were purified from cell debris by sequential centrifugation at 300g (10 min), then 1000g (20 min), and 10,000g (30 min), and filtered through 0.2 μm syringes. Next, the EVs pellet was obtained by centrifugation at 100,000g (60 min), and washed with phosphate buffer solution (PBS). Of note, FBS was depleted from the FBS-derived EVs by centrifugation at 100,000g (2h) prior to the addition to the parent cells. Bradford assay and Nanoparticle Tracking Analysis (NTA) were used to estimate amount of recovered EVs [19].

### 2.5. Characterization of EVs by Nanoparticle Tracking Analysis (NTA), and Atomic Force Microscopy (AFM)

EVs were collected from GDNF-transfected macrophages media, and the size and number of particles were evaluated using NanoSight 500, Version 2.2 (Wiltshire, United Kingdom). In addition, the concentration, and zeta potential (ZP) of EVs were measured using ZetaView QUATT Nanoparticle Tracking Microscope PMX-420 (Particle Metrix, Germany). For this purpose, EVs were diluted to concentration about 2·10^7^ particles/mL with 20 nm filtered PBS. Measurements were performed at 11 positions using the following settings: maximum area 1000, minimum area 5, minimum brightness 20; camera level 16, threshold level 5. Total protein was calculated using standard BCA assay. The AFM imaging was operated as described earlier [42]. A drop of isolated EVs in in 50 mM phosphate buffer, pH 7.4 at total protein 10 µg/mL placed on a glass slide and dried under an argon flow.

### 2.6. Characterization of EV-specific proteins by Western Blot and Label Free Targeted Quantitative Proteomics

The levels of proteins constitutively expressed in EVs (CD9, CD63, CD81, TSG101, and HSP90) were identified by Western blot analysis, using Wes^™^ (ProteinSimple, San Jose, CA, USA). EVs were lysed with 1x RIPA buffer for 30 minutes at room temperature and 200 or 40 mg/ml of protein was denatured and loaded in Wes™ multi-well plates following manufacturer’s instructions. Protein concentrations were determined using BCA kit (Pierce Biotechnology, Rockford, IL, USA). For analysis of CD9, CD63 and CD81 sample’s lysates de-glycosylated using PNGase F PRIME (Bulldog Bio, NZPP050) under non-denaturing conditions at the ratio of 1:9 v:v for 1 h prior to denaturalization. The protein bands were detected with primary antibodies described in **Supplementary Table S2**, and secondary Goat Anti-Rabbit HRP Conjugate (ready-to-use reagent, ProteinSimple, San Jose, CA, USA). A quantitative analysis of obtained images was carried out using Compass SW software.

To characterize EV-GDNF by label free targeted quantitative proteomics, the samples of EVs released by sham-transfected and GDNF-transfected macrophages were digested with trypsin, cleaned up by SPE and prepared for nanoLC-MS/MS analysis as described previously [43, 44] with minor modification. For digestion, to 20 µg of solvent evaporated EV protein was added 50 mM ammonium bicarbonate (100 µL), 40 mM dithiothreitol (10 µL), 0.5 mg/mL β-casein solution (10 µL) (as an indicator of successful digestion and to aid with chromatography retention time verification) and 13.3 µL of 10% sodium deoxycholate (to help with solubilization and denaturation). Samples were denatured for 40 min at 60 °C, shaking at 500rpm, in an Isotemp Thermal Mixer (Fisher Scientific). After cooling, 10 µL of 135 mM iodoacetamide was added and the samples were incubated in the dark at room temperature for 30 min. Ten microliters (10 µL) of a solution containing 1 pmol of an Na^+^/K^+^-ATPase (membrane marker) stable isotope labeled (SIL) peptide standard (purchased from JPT Peptide Technologies, Berlin, Germany) in ∼20/80 acetonitrile/50 mM ammonium bicarbonate was added to each sample. Trypsin (10 µL of 0.1 µg/µL solution in 50 mM acetic acid) was then added to give a 1/20 (w/w) trypsin/protein ratio. Samples were vortexed and digested at 37 °C for 20 h, shaking at 300 rpm, in the Isotemp Thermal Mixer. After digestion, 10 % TFA solution was added to stop the reaction, such that the volume added was 10 % of the total volume of the digestion reaction. A deoxycholate precipitate formed. The precipitate was pelleted by centrifuging at 13.4K x *g* (5 min). The obtained samples were purified on the SPE cartridges with polymeric reversed phase, 10 mg/mL, and eluted into LoBind Eppendorf tubes with 60% acetonitrile/40% formic acid. Then, the evaporated solution was reconstituted with 2% acetonitrile. The reconstituted sample was vortexed and then centrifuged at 13.4K x *g* for 5 min. The supernatant was transferred to deactivated vial inserts (part # WAT094171DV; Waters, Milford, MA) before nanoLCMS/MS analysis.

The nanoLC-MS/MS analysis was performed on a nanoAcquity UPLC® (Waters) coupled to a SCIEX QTRAP 5500 (Framingham, MA) hybrid mass spectrometer with NanoSpray® III source. The specifics of this analysis have been previously reported in [23, 43].

The sequences of peptides that could be identified and were used in the final quantitative assessment are shown in **Supplementary Table S3**. UniProt accession numbers and the MRMs employed for each peptide have been previously published [23]. The test samples were analyzed in duplicate in the same batch, the method being both label and standard free.

### 2.7. Characterization of EV-GDNF by quantitative qPCR Analysis

EVs or cells were lysed and directly added to qPCR reactions. qPCR was performed on lysates with 0.5 μL each of 20 μM forward and reverse primers 0.5-2 μL of sample and 6.25 μL PowerUp SYBR Green Master Mix (Applied Biosystems) with a total reaction volume of 12.5 μL using a QuantStudio 6 Flex Real-Time PCR System (Applied Biosystems). Gene specific primers were used to amplify GDNF (forward sequence 5-GCAGACCCATCGCCTTTGAT-3 and reverse 5-CCACACCTTTTAGCGGAATGC-3).

### 2.8. Animals

Parkin Q311X(A) mice (two breeding pairs, 12 weeks old) were obtained from the Jackson Laboratory (Bar Harbor, ME, USA) and used to start a colony. The animals were treated in accordance with the Principles of Animal Care outlined by National Institutes of Health and approved by the Institutional Animal Care and Use Committee of the University of North Carolina at Chapel Hill; “Inflammatory cells for transport of therapeutic polypeptides across the Blood Brain Barrier” ID# 18-064.0, Web ID: 644959, date for renewal 02/28/2021. PCR analysis was used for identification of transgenic mice and wild type mice as described earlier [38]. All experiments were performed in 4 – 16 mo. old male Parkin Q311X(A) mice on a C57BL/6 genetic background with age-matched non-transgenic littermates served as controls. Animals were housed in a temperature and humidity-controlled facility on a 12 h light/dark cycle and food and water were provided ad libitum. For all experiments, mice were monitored for any adverse signs of discomfort.

### 2.9. Treatment of animals

Four groups of mice (*N* = 10) were employed in these studies. Three groups of Parkin Q311(X)A mice (4 mo. old) were intranasally injected with EV-GDNF (once a week, three weeks, 3 × 10^9^ particles/10 uL/mouse), or sham EVs (the same number of particles), or buffered saline (0.9% sodium chloride, pH = 7.4, negative control). Another froup with wild type (WT) mice was injected with saline and used as a positive control.

For intranasal injections (*i.n*.), mice were anesthetized with isoflurane (2 % during induction, 1.5% - 2% during maintenance), until it showed no signs of reaction on four paw toe pinch. The ample sterile ophthalmic gels were applied to ensure the moist state of corneas of the animal. Then, the mouse was placed on a clean drape facing up with a heating pad underneath to maintain body temperature. A padded pillow made of rolled up paper towels with tapes is adjusted to ensure the upright angle of the nostrils when placed under the head of the mouse. Using a micropipette, 5 μL of a treatment solution formulation was dispensed into each nostril of the mouse. The aspiration of the droplet was visually confirmed before moving to the next mouse. Animals were allowed to regain mobility in a recovery chamber, with the supine position maintained throughout. Mice were observed for recovery from sedation during 30 min after the drug *i.n*. administration. In particular, breathing, movement, and overall being were be monitored.

### 2.10. Behavioral Studies

All four animal groups were subjected to standard behavioral tests before the treatment and later on for one year. To evaluate therapeutic effects of EV-GDNF on locomotor activity and rearing movements, a Wire hanging task for grip strength, an accelerating Rotarod procedure, and an open field test (OFT) were performed. *<u>Wire hanging test for grip strength</u>*. Each mouse was placed on a wire, and latency for the mouse to fall from the wire was recorded. The maximum trial length was 180 sec. *<u>Rotarod test</u>*. For the traditional constant speed Rotarod test, mice were trained and tested as previously described with slight modifications [45]. The accelerating rotarod (Ugo Basil) was used for assessing motor coordination, balance, and ataxia. A rotarod machine with automatic timers and falling sensors was used. The mouse was placed on a 9 cm diameter drum. The surface of the drum was covered with hard chloroethylene, which does not permit gripping on the surface. Before the training sessions, the mice were habituated to stay on the stationary drum for 3 min. Habituation was repeated every day for 1 min just before the session. Mice were placed on a cylinder, which slowly accelerates to a constant rotating speed. Normal mice readily learn to walk forward as the drum turns. For each trial, revolutions per minute (rpm) are set at an initial value of 5, with a progressive increase to a maximum of 30 rpm across 5 min, the maximum trial length. For the first session, mice were given 3 trials, with 45 sec between each trial. Measures were taken of latency to fall from the top of the rotating barrel. A second test session with 2 trials is conducted 48 h later, to evaluate consolidation of motor learning. *<u>Activity in an open field</u>*. The hyperactivity of PD mice treated with different EV-based formulations, as well as WT mice injected with saline as controls (*N* = 10) was assessed by five-minutes and one-hour trials in an open field chamber (41 cm x 41 cm x 30 cm), crossed by a grid of photobeams. Measures were taken by an observer blind to mouse genotype and treatment. Counts were taken of the number of photobeams broken during the trial, with separate measures for vertical rearing movements (VersaMax, AccuScan Instruments). Time spent in the center regions was used as an index of anxiety-like behavior. Behavioral data were analyzed using one-way or repeated measures Analysis of Variance (ANOVA). For all comparisons, significance was set at *p* < 0.05.

### 2.11. Immunohistochemical and Stereological Analyses

At the endpoint (16 mo. old) animals were sacrificed, and perfused. Postmortem brains were harvested, washed, postfixed, and immunohistochemical analysis was performed in 30 µm thick consecutive coronal brain sections. Two methods of sectioning the tissues were utilized. For investigations of the effect of EV-formulations on the protection of dopaminergic neurons, as well as decreasesn in microglial activation, the free-floating sections of the tissues were used. This approach allows to see the structure of dendrites and axons through the sample. The sections were not mounted on slides until after completing the immunohistochemistry process. To examine *<u>neuroprotective effects</u>* of EV-based formulations, dopaminergic neurons were visualized with tyrosine hydroxylase staining as described in [46]. To study *<u>microglial activation</u>* in the brain of treated animals, primary monoclonal rat anti mouse Mac1 antibodies (AbD Serotec, Raleigh, NC, USA) were used along with secondary biotinylated goat anti-rat antibodies (Vector Laboratories, Burlingame, CA, USA). Each midbrain section was viewed at low power (10x objective), and the *SNpc* was outlined [47].

For the visualization of neurons, brain sections were stained with cresyl violet acetate solution (*<u>Nissl staining)</u>*. For this purpose, mice were sacrificed, perfused, and brains collected were frozen in optimal cutting temperature media (OCT). Frozen brains were cut into coronal sections at 10 µm thickness using a CryoStarTM NX50 Cryostat (ThermoFisher) and then placed on a warm pad to remove excess OCT. Slides were stored at -20°C. A subset of sliced tissues was stained with Cresyl violet acetate solution (Nissl) for the detection of Nissl body in the cytoplasm of neurons that stained purple blue. In brief, tissues were exposed to xylene and rehydrated in a graded series of ethanol at 100, 95 and 75% concentrations. Sections were exposed with Nissl staining solution for 15 min, washed in distilled water, immersed in ethanol at 75%, 95%, and 100% concentrations, and then cleared by xylene. Slides were then mounted using mounting media for visualization. Three to five areas were randomly selected to be examined with an inverted fluorescence microscope and 63X objectives (Zeiss, Germany) by investigators who were blinded to the experimental groups. Another subset of sliced tissues was stained with *<u>Hematoxylin and Eosin (H&E)</u>* for the detection of morphological changes within the different brain regions. Tissues were exposed to xylene and rehydrated with ethanol, at 100, 95 and 70% concentrations, followed by staining with hematoxylin dye for 15 minutes. After washes in distilled water, tissue sections were stained with eosin for 20 seconds and then dehydrated with gradient ethanol. Sections were exposed to xylene and then mounted using mounting media for visualization. Sections were imaged as described above.

### 2.12. ELISA

To examine potential antiinflammatory effects of EV-GDNF formulations in mice, the levels of cytokines, Interferon γ (IFN-γ), Interleukin 4 (IL-4), Interleukin 6 (IL-6), Interferon-inducible Protein 10 (IP-10), and tumor necrosis factor alpha (TNF-α), and chemokines, Monocyte Chemotactic Protein-1 (MCP1), regulated on activation, normal T cell expressed and secreted (RANTES), were analyzed in the spleen, brain, and liver recovered at necropsy. For this purpose, organs were removed post-mortem and homogenized in cell lysis buffer using an electrical homogenizer apparatus. Supernatant was used to measure secretion levels of the inflammatory molecules by ELIZA (R&D Systems) according to the manufacturer’s instructions. The optical density was read at A450 on a Synergy HTX plate reader. Results are shown as the mean ± the standard error of the mean (SEM).

### 2.13. Statistical Analysis

For all experiments, data are presented as the mean ± SEM. Tests for significant differences between the groups in experiments regarding *characterization of EVs* released by different types of parental cells were performed using a one-way ANOVA with multiple comparisons (Fisher’s pairwise comparisons) using GraphPad Prism 5.0 or higher (GraphPad software, San Diego, CA, USA). For targeted quantitative proteomics *data*, the MRM peak area data was processed with MultiQuant 2.0.2 software (SCIEX). Peak areas for the three highest responding MRMs for each peptide were summed and responses between samples were compared for each peptide. The presence of multiple peptides for a protein provided extra confidence that the protein/tetraspanin/integrin was present. SIL or label free standards would be needed to compare abundance of proteins within a sample, large differences in signal sometimes being seen between peptides of the same amount. T-tests were used to determine whether peptide abundance differences between samples were significant (*p* < 0.05). For *in vivo* experiments, 10 mice were used per group. Thus, for behavioral tests the total number of mice was determined by power analysis: rotarod tests and open field tests of therapeutic effect of EV-mediated GDNF delivery from previously obtained by us data, wherein a = 0.05, s = 1640, and power = 0.90. To allow equal sample sizes/group for ANOVA analysis, the number of mice necessary will be 10 recipient mice/treatment group. This related to variability of inflammatory responses in mice. Next, for a histological evaluation of tyrosine hydroxylase positive neurons, numbers of recipients were determined from retrospective power analysis using the SAS JMP program. For dopaminergic loss, a 30 % loss and subsequent neuroprotection from dopaminergic loss for = 0.05, and = 2251 with a power of 0.80 require 10 recipient mice/treatment group. For inflammatory responses, for example, 30 % diminution of inflammatory response by number of Mac-1+ reactive microglia/section for = 0.05, and = 4.6 with a power of 0.80 require 10 recipient mice/treatment group. For the evaluation of pro-inflammatory cytokines levels in PD mice, results were shown as the mean ± the standard error of the mean (SEM). For multiple factors a two-way ANOVA followed by post-hoc tests as appropriate (Tukey or Dunnett’s) were used for multiple comparisons using Prism 8.0 (GraphPad Software). *p* < 0.05 was considered statistically significant.

## 3. Results

### 3.1. Manufacture of EV-GDNF by Genetic Modification of Primary Macrophages

In this work, we utilized endogenous loading of EVs through the parent cells. Based on our previous investigations [23, 27, 47-49], primary macrophages were selected for production of EV-GDNF formulation. Briefly, macrophages were transfected with GDNF-encoding plasmid DNA (*p*DNA) by electroporation at different electroporation conditions (as described in **Materials and Methods** section), and the levels of GDNF in parent cells and in EVs collected from macrophage conditioned media were assessed by ELISA over six days after the transfection (**Supplementary Figure S2**). Significant levels of GDNF were detected in transfected macrophages (solid bars), as well as EVs isolated from the media (striped bars). Of note, the GDNF expression levels in macrophages gradually decreased from day 1 to day 6, although, the amount of GDNF in EVs increased, especially at later days. The variation of electroporation conditions (#2 - #4) did not significantly affected the GDNF amount in EVs, so the condition #4, which provided slightly higher GDNF content in EVs, was selected for further investigations.

### 3.2. Characterization of EV-GDNF

EV-GDNF collected from genetically modified parent macrophages were characterized for size, charge, shape, and morphology. Sham EVs collected from sham transfected macrophages were used as a control. As seen on **Figure 1**, the transfection of parent cells with GDNF-encoding *p*DNA did not significantly affect size and charge of the nanocarriers. According to NTA data, average mode size 120 nm with negative ZP around -20 mV was recorded for both EV-GDNF and sham EVs (**Figure 1 A**). Next, the spherical morphology of the obtained EV-GDNF with relatively uniformly size distribution was confirmed by AFM (**Figure 1 B**). Finally, the presence of EV-specific proteins (CD9, CD63, CD81, TSG101, and SHP90) in EV-GDNF was confirmed by Wes™ Simple Western Blot **(Figure 1 C)** and quantified using Compass SW software (**Figure 1 D**).

**Figure 1.**
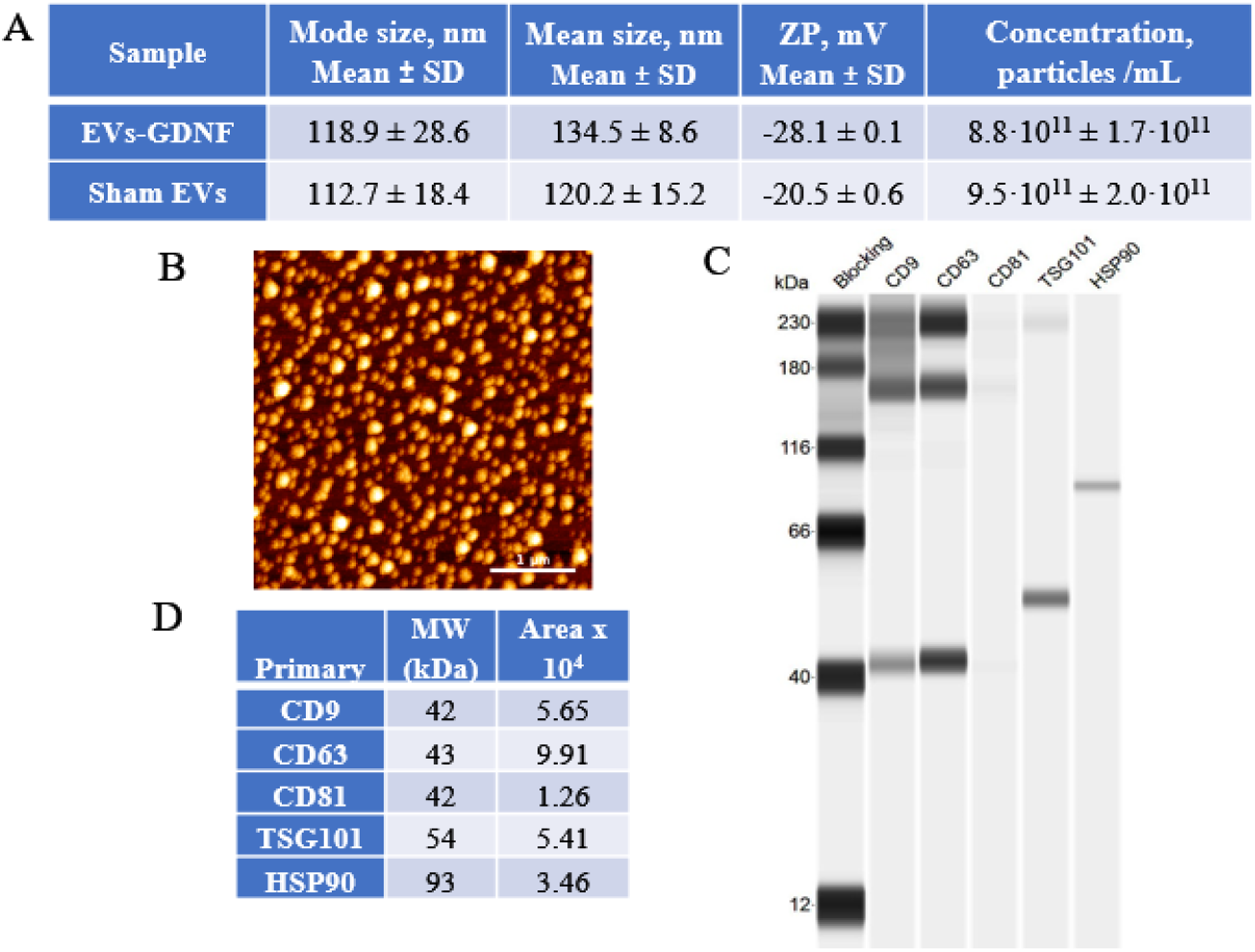
Characterization of EV-GDNF by ZetaView QUATT Nanoparticle Tracking Microscope PMX-420, AFM, and Western blot. Primary macrophages were transfected with GDNF-encoding *p*DNA by electroporation (condition #4), and EV-GDNF were collected from conditioned media on day 6. EV-GDNF were characterized for size, zeta potential, and morphology by ZetaView QUATT Nanoparticle Tracking Microscope PMX-420 (**A**), and AFM (**B**). The presence of EV-specific membrane proteins was EV-GDNF was confirmed by Wes (**C**) and quantified using Compass SW software (**D**). The bar: 1 µm.

We reported earlier that similar to their parent cells, macrophage-derived EVs exert unique biological activity reflective of their origin that allows them easily penetrate the BBB and migrate rapidly to sites of neurodegeneration [50]. It is of paramount importance in the view that chronic brain inflammation is a common feature for different neurodegenerative disorders, including PD patients [51]. In addition, EVs has different adhesive proteins expreseed on their membranes that allow efficient binding and delivery of their cargo to target cells [52]. However, genetic modification of parent macrophages may alter their composition. Therefore, we investigated the expression of the EV-specific proteins in EV-GDNF by label free targeted quantitative proteomics. Specifically, we assesed the relative expression levels of adhesive tetraspanins (CD63 and CD9), ALIX, integrin β-1 (CD29), and tsg101 (**Supplementary Table S3**) released by GDNF-transfected macrophages and compared with those released by sham-transfected cells. Additionally, we studied the presence of α Integrins and Integrin β-2 that facilitate targeting to the inflamed endothelium. The data revealed that transfection of parent macrophages did not statistically significantly affect expression of the proteins of interest (**Supplementary Figure S3**), suggesting that similar to sham EVs, EV-GDNF could successfully deliver therapeutic cargo to the inflamed brain.

Finally, the presence of GDNF-encoding DNA in EVs released by GDNF-transfected parent cells was studied by qPCR analysis (**Figure 2**). The obtained data suggested that these EVs contained significant amount of GDNF-DNA. This result is consistent with our previous reports [21, 53] indicating that EVs secreted from GFP- and TPP1-transfected parent macrophages contain corresponding genetic material.

**Figure 2.**
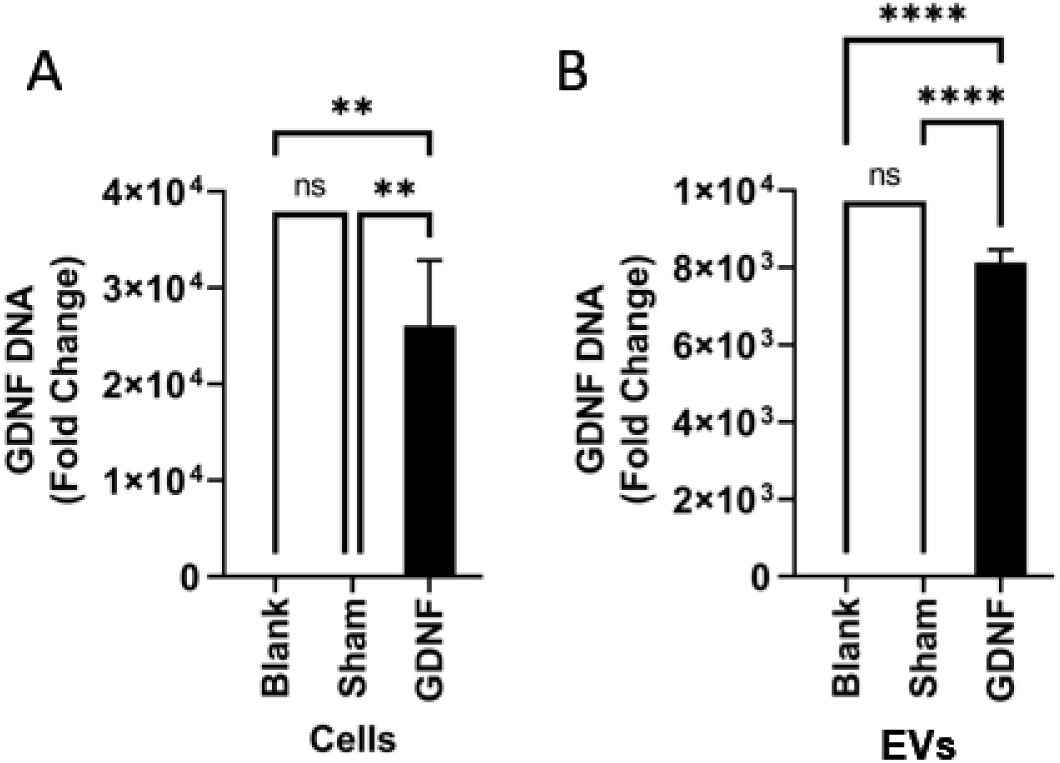
Characterization of genetic content of GDNF-transfected parent macrophages and released EV-GDNF by quantitative qPCR. Macrophages were transfected with GDNF-encoding *p*DNA by electroporation, and the levels of GDNF-DNA in the cells (**A**) and EVs released by these cells (**B**) were assessed. A significant amount of GDNF-DNA was detected in parent cells, as well as in the EVs. Statistical significance was assessed by One Way ANOVA corrected for multiple comparisons using the FDR. \*\**p* < 0.01, or \*\*\*\**p* < 0.0001.

### 3.3. Effect of EV-GDNF on Locomotor Activity upon Intranasal Administration to Parkin Q311(X)A mice

Based on our previous reports [19], we selected intranasal (*i.n*.) administration route for treatment with EV-based formulations, as one of the most efficient routes for the brain delivery. Transgenic PD mice (4 mo. old) were treated with EV-GDNF (3 × 10^9^ particles/10 µL/mouse) once a week three times, and their locomotor functions were assessed in a battery of behavioral tests (**Figure 3**). PD mice and wild type (WT) counterparts treated with saline were used as positive and negative controls, respectively. PD mice injected with sham EVs were used in another control group. The behavioral tests including Wire hanging test (**Figure 3A**), and Rotarod test (**Figure 3B**) were performed for all animal groups over a year.

**Figure 3.**
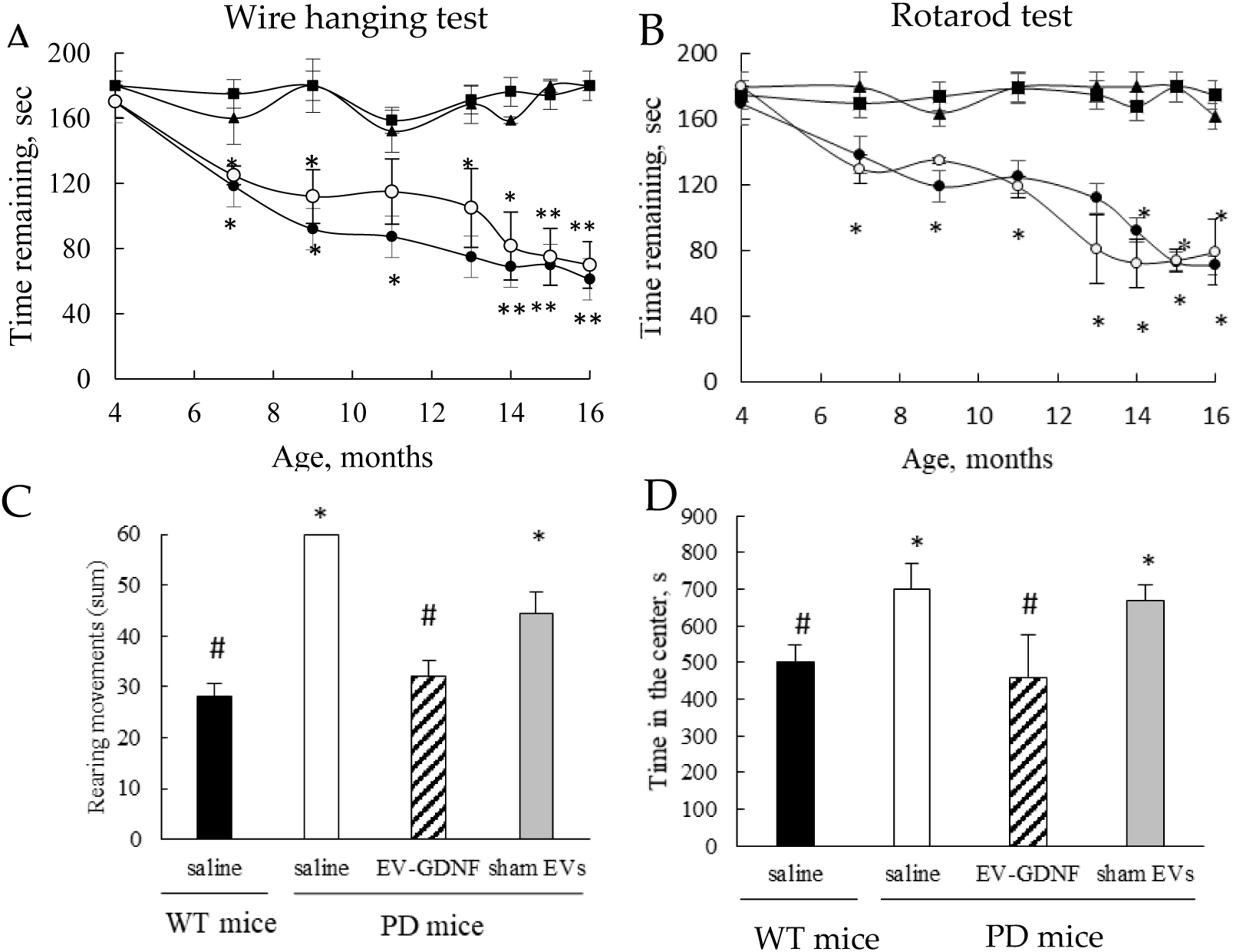
Behavioral tests demonstrating significant therapeutic effect of EV-GDNF in Parkin-Q311X(A) mice. The effect of EV-GDNF on motor functions and activity was assessed in Wire hanging test, and Rotarod test (**A, B**), as well as in OFA tests (**C, D**). (**A, B**) Transgenic mice were *i.n*. injected with EV-GDNF (triangles, 3×10^9^ particles/10 µL/mouse), or sham EVs (empty circles, 3×10^9^ particles/10 µL/mouse), or saline (filled circles, 10 µL/mouse). Wild type mice were *i.n*. injected with saline (filled squares, 10 µL/mouse) were used as controls. Wire hanging test (**A**), and Rotarod test (**B**) demonstrated significant improvements in motor functions upon treatment with EV-GDNF. (**C, D**) OFA tests at 12 mo. demonstrated improved behavior in EV-GDNF treated PD mice (striped bars) compared to PD mice treated with saline (white bars) that was similar as in healthy WT mice (black bars) including decreases in the hyperactivity and anxiety-like behavior. The differences between sham EVs and saline in PD mice were inconclusive. Values are means ± SEM (*N* = 10), * *p* < 0.05, ** *p* < 0.005, and #*p*< 0.05, as compared to WT control.

The obtained data indicated significant therapeutic efficacy of EV-GDNF treatments reflected in preservation of locomotor functions in PD mice (**Figures 3A, B**; triangles) compared to PD mice treated with saline (**Figures 3A, B**; filled circles). Moreover, the remaining time in Wire hanging test, and latency to fall in Rotarod test were almost the same as in WT healthy animal group (**Figures 3A, B**; squares). Of note, the preservation of locomotor activity in PD mice by EV-GDNF was recorded as long as for a year, and was almost the same as in healthy WT animals (**Figures 3A, B**; filled squares). No statistically significant effects were detected in the transgenic mice treated with sham EVs (**Figures 3A, B**; empty circles), indicating these comparisons were inconclusive.

To reinforce this conclusion, we performed Open Field Activity (OFA) tests with 16 mo. old animals (**Figure 3 C, D**). A one-hour trial was completed in an OF chamber (41 cm x 41 cm x 30 cm) equipped with crossing grid of photobeams (VersaMax system, AccuScan Instruments) to assess the effect of therapeutic treatments on hyperactivity of PD mice. The number of rearing movements of the animals, and the amount of time spent in the center of the chamber, a known index of anxiety-like behavior, were recorded. The OFA tests indicated that PD mice treated with saline (white bars) showed hyperactivity with more errors per step while traversing the beam (**Figure 3 C**), and anxiety-like behavior manifested in more time spent in the center region (**Figure 3 D**) compared to WT mice treated with saline (black bars). In contrast, PD mice treated with EV-GDNF (striped bars) displayed improved behavior that was similar as in healthy WT mice treated with saline. Whether the treatment with sham EVs (grey bars) affected the performance patterns of PD mice was inconclusive, and the PD mice demonstrated similar hyperactivity and anxiety-like behavior as control PD animals treated with saline.

### 3.4. Neuroprotective and Anti-inflammatory Effects in ParkinQ311(X)A mice

One year after the first treatment, mice (16 mo. old) were sacrificed, brains were isolated, and sections were mounted on slides. The mid-brain slides were stained for the expression of tyrosine hydroxylase (TH), a marker for dopaminergic (DA) neurons (**Figure 4 A**). Almost complete degeneration of DA neurons in the *SNpc* was observed in PD mice treated with saline compared to healthy WT mice. In contrast, treatments with EV-GDNF dramatically ameliorated PD-related neurodegeneration. Furthermore, potent anti-inflammatory effects of EV-GDNF was demonstrated (**Figure 4 B**). Thus, the neurodegeneration in 16 mo. of age Parkin Q311(X)A mice treated with saline was accompanied with substantial brain inflammation as displayed by up-regulated expression of CD11b by microglia within the *SNpc* that exhibited a more amoeboid morphology with larger cell body, compared to WT mice treated with saline. Importantly, the treatment with EV-GDNF significantly reduced neuroinflammation in PD mice according to decreased microgliosis revealed by ramified microglia. No anti-inflammatory effects of sham EVs was found in PD mice.

**Figure 4.**
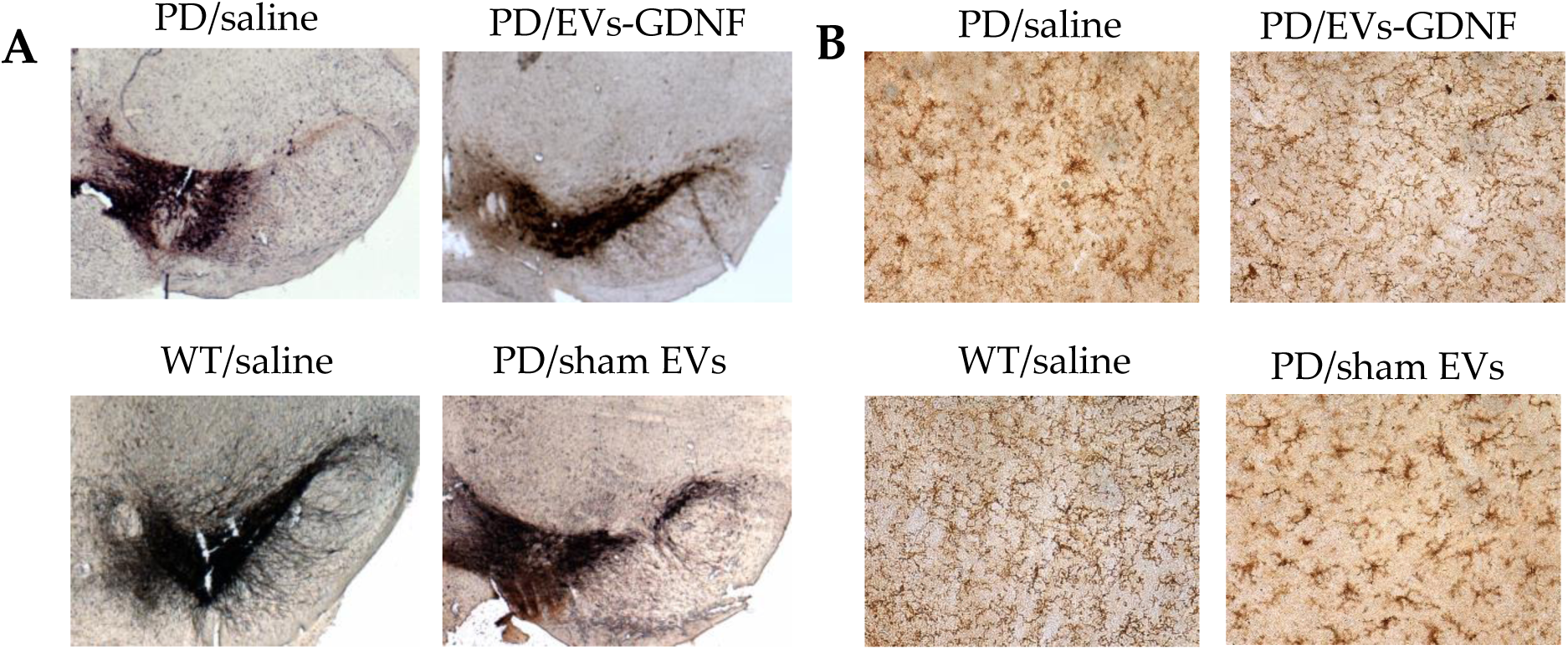
Neuroprotective and antiinflammatory effects of EV-GDNF in Parkin Q311(X)A mice. Transgenic mice (4 mo. old, *N* = 10) were *i.n*. injected with: saline (10 µL/mouse), or EV-GDNF (3×10^9^ particles/10 µL/mouse), or sham EVs (3×10^9^ particles/10 µL/mouse). Wild type control mice were intranasally injected with saline (10 µL/mouse). Animals were sacrificed at mo. 16, and brain slides were stained with TH, a marker for dopaminergic neurons (**A**); or Ab to CD11b for activated microglia (**B**). The images indicate significant preservation of TH-positive neurons and decrease in microglial activation in Parkin Q311(X)A mice upon EV-GDNF treatment compared to PD mice treated with saline. The administration of sham EVs did not cause significant therapeutic effects.

Furthermore, the antiinflammatory effects of EV-GDNF treatments were confirmed by measuring the levels of pro-inflammatory cytokines and chemokines in main organs of animals (**Figure 5**). Significant (#) secretion of IFN-γ, IL-6, MCP-1 and TNF-α were identified in the brain of PD mice injected with saline and of IL-6, IP-10 and TNF-α in sham EVs, when compared to healthy control. In contrast, PD mice treated with EV-GDNF showed a significant decrease (*) in secretion of IFN-γ and MCP-1 when compared to PD mice treated with saline, and a significant decrease ($) in IL-4, IL-6, IP-10 and TNF-α, when compared to PD mice treated with saline and sham EVs, similar to WT mice, or even lower. This confirms that EVs loaded with GDNF could diminish inflammation in the brain of PD mice.

**Figure 5.**
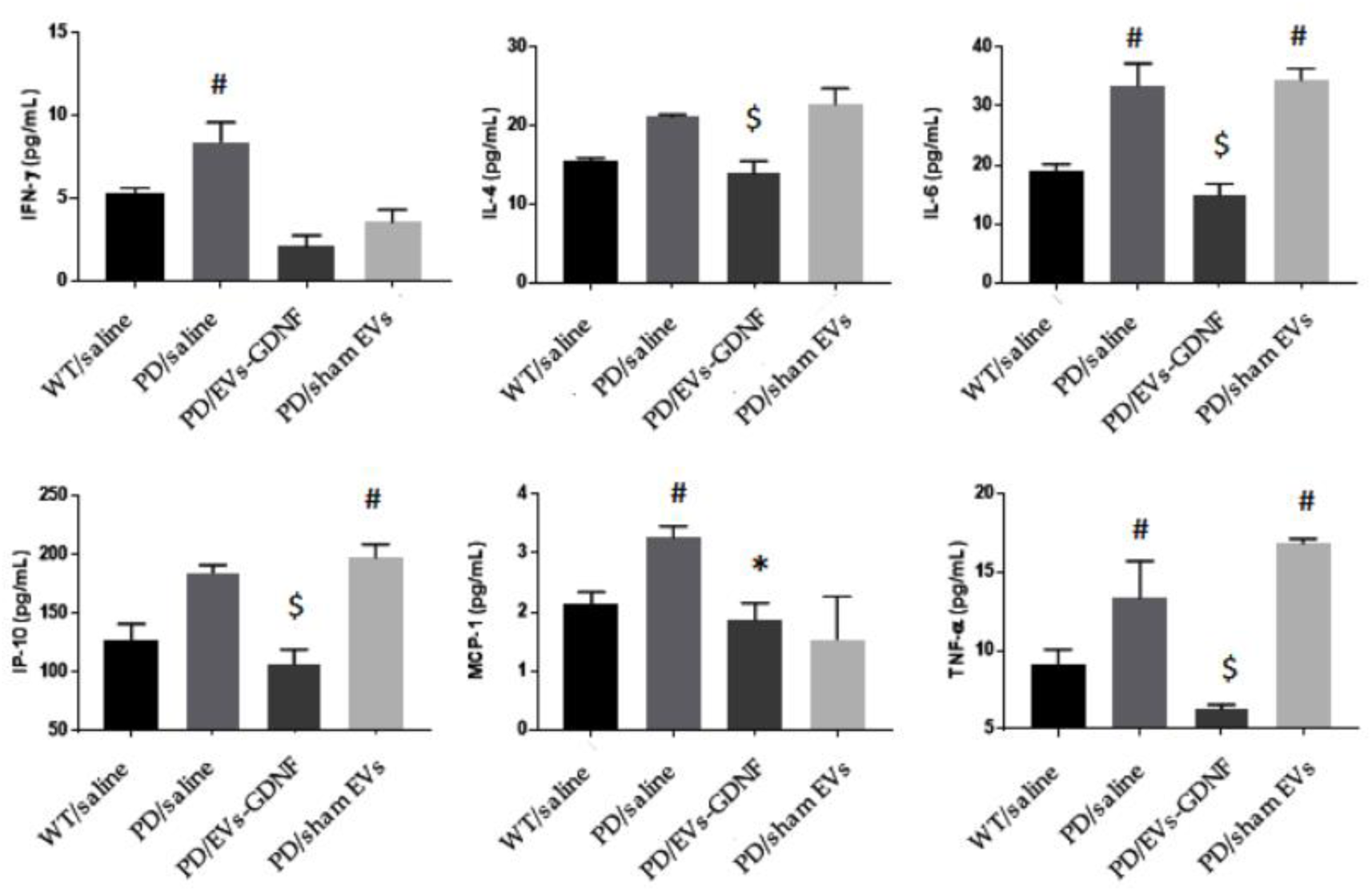
Anti-inflammatory effects of GDNF-carrying EVs in PD mouse model. Transgenic mice (4 mo. old) were intranasally injected with: saline (10 µL/mouse), or EV-GDNF (3×10^9^ particles/10 µL/mouse), or sham EVs (3×10^9^ particles/10 µL/mouse). Wild type control mice were intranasally injected with saline (10 µL/mouse). Animals were sacrificed at mo. 16, brains were removed post-mortem, and homogenized in cell lysis buffer. Elevated cytokine levels in the brain, were recorded in PD mice treated with saline and Sham EVs. Administration of EV-GDNF significantly decreased pro-inflammatory molecules in the brain compared with PD mice treated with saline. *N* = 4, #*p* < 0.05 compared to healthy WT animals; \**p* < 0.05 compared to PD mice treated with saline, ^$^*p* <0.05 compared to PD mice treated with saline and sham EVs.

The neuroprotective effects of EV-based formulation in Parkin Q311(X)A mice were further confirmed with the Nissl staining (**Figure 6 A-D**). While a large quantity of healthy neurons was found in the *SNpc* of WT mice (**Figure 6 A**), PD mice treated with saline showed lower number of neurons and astrocytes (**Figure 6 B**). In contrast, brains of PD mice treated with EV-GDNF showed better tissue integrity, with minimal signs of vacuolation in neurons (**Figure 6 C**). Of note, treatments with sham EVs did not have this therapeutic effect (**Figure 6 D**). The same effect of neuronal protection in PD mice by EV-GDNF was shown by H&E staining (**Figure 6 E-H**). In particular, healthy cell morphology and low vacuolation was detected in WT mice (**Figure 6 E)**, while damaged tissue with degeneration in the neurons (black arrows) and elongated irregular nuclear morphology of microglia (blue arrow) were observed in brain slides of PD animals treated with saline (**Figure 6 F**). Moreover, the vacuolation within the overlying molecular layer suggests potential swelling/degeneration of Purkinje neuron dendrites. Healthier looking tissues with high integrity of neurons were visible in tissues of mice treated with EV-GDNF (**Figure 6 G**) indicated neuroprotection, compared to PD mice treated with saline (**Figure 5 G**). However, elongated nuclear morphology of microglia (blue arrow) and some necrotic neurons (black arrows) were detected albeit noticebly less than in PD animals triated with saline (**Figure 6 F**). In accordance with all other assays, sham EVs did not produce significant therapeutic effects in PD mice (**Figure 6 D** and **H**). Additional images of brain slides with Nissil staining and H&E staining are presented on Supplementary Figures S4 and S5, respectively.

**Figure 6.**
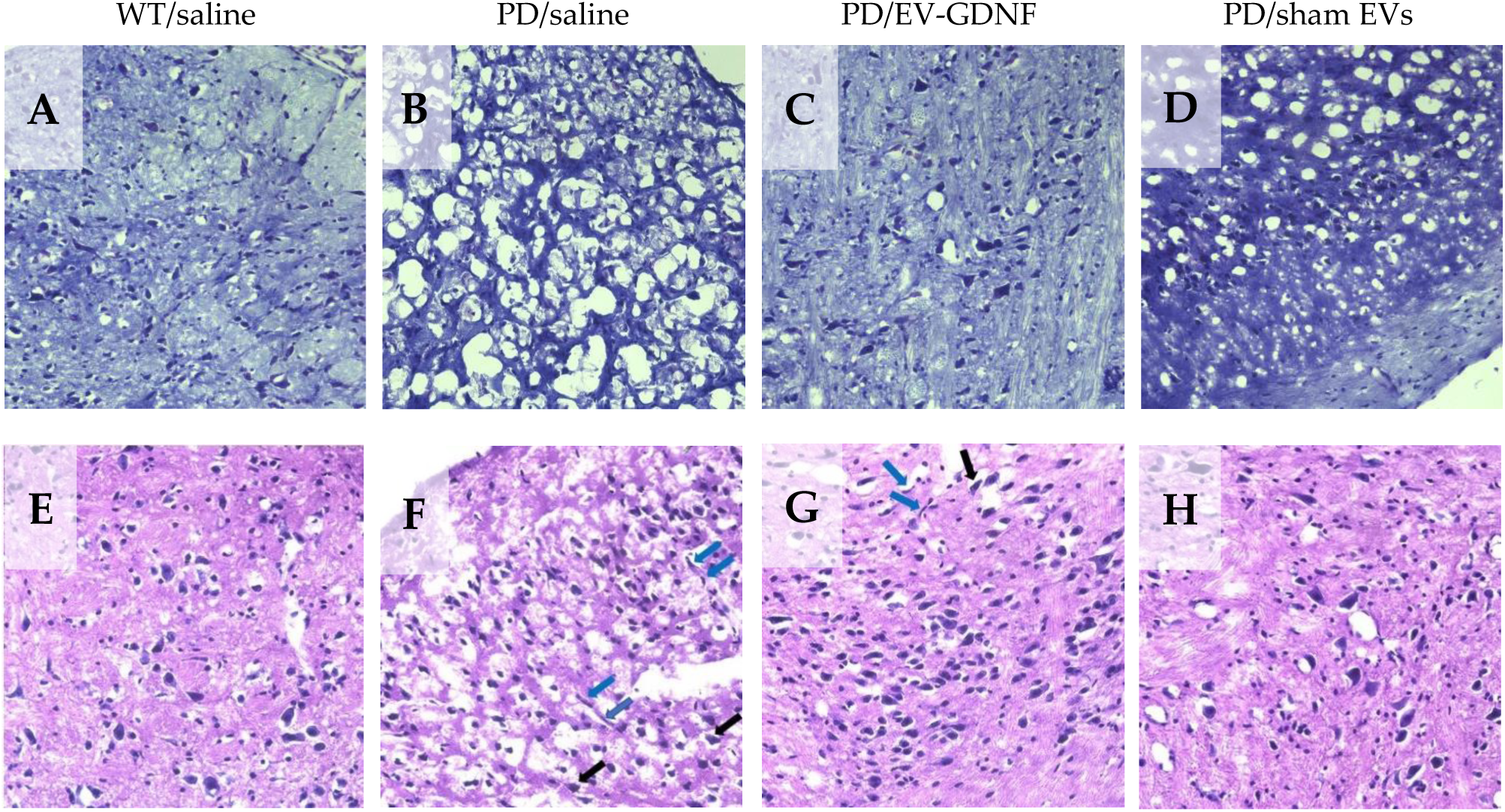
Histological analysis of neuroprotective effects by EV-GDNF in Parkin Q311(X)A mice. Transgenic mice (4 mo. old) were intranasally injected with: saline (10 µL/mouse), or (**3**) EV-GDNF (3×10^9^ particles/10 µL/mouse), or sham EVs (3×10^9^ particles/10 µL/mouse). Wild type control mice were intranasally injected with saline (10 µL/mouse). Animals were sacrificed at mo. 16, brain slides were stained with Nissl staining (**A** – **D**) and H&E staining (**E** –**H**). The obtained bright light images show lower number of Nissl bodies with neuronal shrinkage (**B**) and damages tissues with degeneration in the neurons (**F**) in PD mice treated with saline when compared to WT mice (**A, E**). Histological analysis indicate neuroprotective effects in the brain of PD mice treated with EV-GDNF with healthy morphology in tissue structure and high integrity of neurons (**C, G**) when comparted to PD mice treated with saline (**B, F**). The administration of sham EVs did not have significant therapeutic effect in PD mice (**D, H**). Black arrows, degenerated neurons; blue arrows, elongated irregular nuclear morphology.

Finally, possible inflammatory effects of EV-GDNF formulation were also examined in the main peripheral organs, liver and spleen. We demonstrated earlier that these organs accumulate the most amounts of EVs in mice [21, 23]. For this purpose, levels of pro-inflammatory cytokines, IFN-γ, IP-10, IL-4, IL-6, RANTES, MCP-1, and TNF-α were assesed in PD mice treated with EV-GDNF, and compared with those in WT animals treated with saline (**Supplementary Table S4**). No significant increases in cytokine levels were found followed administration of EV-GDNF. This is consistent with our previous reports indicated no significant toxicity of EVs [54]. Furthermore, no total weight loss was detected in mice treated with EV-GDNF, or sham EVs compared to those treated with saline (**Supplementary Figure S6**). This signifies absence of general toxicity of the cell-based formulations upon multiple administrations of EVs treatments.

## 4. Discussion

In this work, we exploited macrophage derived EVs as promising bio-nanostructures that allow targeting disease tissues and efficient delivery of incorporated therapeutics to the brain. We posit that this approach has a potential to become a novel treatment strategy that links a cutting-edge class of precision medicines with the latest nanotechnology. We reported earlier development of EV-based therapeutic formulations using exogenous loading of naïve EVs with a number of potent therapeutic proteins, including antioxidant enzyme, catalase [19], brain-derived neurotrophic factor (BDNF) [22], and tripeptidyl peptidase-1 (TPP1), a therapeutic enzyme for treatment of a lysosomal storage disorder, Batten disease [21]. However, in some cases therapeutic proteins may be not available in significant amounts for exogenous loading into EVs. Furthermore, many therapeutic molecules, especially bioactive proteins, and enzymes, are highly susceptible for the degradation or deactivation upon loading. Herein, we utilized a different strategy, the endogenous loading of GDNF through the transfection of parent cells with GDNF-encoding *p*DNA.

First, we optimized transfection conditions that allowed efficient genetic modification of parent autologous macrophages and manufacture a considerable amount of EVs with prolonged expression of GDNF. We characterized obtained EV-GDNF according to MISEV2018 guidelines for size, charge, shape, morphology, and protein content. Spherical nanoparticles with relatively uniform size around 120 nm were confirmed by NTA analysis, and AFM. The size of the particles appeared to be on the upper side of the typical median particle size to ensure successful deposition within the nasal cavities. Thus, it was reported that larger than 130 nm particles may deposit at the front of the nose, while finer particles tend to penetrate further into the brain tissues [55]. Furthermore, the presence of EV-specific proteins that are known to facilitate EVs adhesion to target cells was verified for EV-GDNF by western blot.

It is of paramount importance to ensure that GDNF was incorporated /associated with EVs in the developed formulation and not just co-isolated with these drug carriers. To this point, we reported earlier characterization of EVs isolated from GDNF-transfected macrophages media by Western blot [40]. We demonstrated that GDNF was protected in EVs against degradation by pronase. At the same time, free GDNF was completely degraded at the same conditions. Destruction of EVs by sonication eliminated this protective effect. This indicates that at least significant portion of GDNF molecules was incorporated into EVs. Nevertheless, we cannot completely state that all GDNF molecules are in the EVs lumen; some portion can be associated with the EVs membrane. Furthermore, we demonstrated here that along with the encoded therapeutic protein (GDNF), EVs released by pre-transfected macrophages contain genetic material, GDNF-DNA. This may result in transfection of brain tissues and GDNF expression at the disease site. We speculated that a specific mechanism that results in DNA targeted accumulation of EVs might exisit in the parent cells. The further investigations regarding specific location of GDNF on EVs and the possibility of transfection of brain tissues by EV-GDNF are ongoing in our lab.

Regarding targeting drug delivery by EVs nanocarriers, it is well established that specialized cells of the immune system, including monocytes, macrophages, and T cells, can accumulate in the PD brain migrating to the sites of neuroinflammation and degeneration [56, 57]. Moreover, this ability to target inflamed tissues was also shown for macrophage derived EVs [19, 21, 36] that is of particular importance due to the fact that pathological process in the brain of PD patients is accompanied with chronic neuroinflammation [51]. Herein, using label free targeted quantitative proteomics, we confirmed that genetic modification of parent macrophages with GDNF-encoding *p*DNA did not significantly alter the expression levels of specific integrins on EVs that promote adhesion and targeting tissues with inflammation. This suggests that the obtained EV-GDNF would accumulate in the PD mouse brain and potentially deliver their therapeutic cargo to the disease site.

For the assessment of therapeutic efficacy of EV-GDNF in transgenic mouse model of PD, we initiated treatments *via* intranasal (*i.n*.) administration at early stages of the disease. This route provides two different paths for transport to the brain. These are: *i*) the passage along the olfactory nerve cells, when nanoparticles bypass the BBB and enter the brain directly, and *ii*) the transport across the epithelial cell layer to the systemic blood circulation, and then transferring across the BBB into the brain parenchyma. Of note, the first route allows entering the CNS without first-pass hepatic and intestinal metabolism that may significantly reduce EVs clearance in these peripheral organs [58]. Thus, the mucosa and lamina propria are exceedingly vascularized with the high absorption rate epithelium [59]. Furthermore, this non-invasive route of drug delivery has a high patient comliancy and does not requre frequent hospital visits. We also demonstrated earlier that considerable amount of EVs administered through *i.n*. route accumulated in the mouse brain with neuroinflammation [19]. Thus, confocal images of PD mouse brain showed diffuse staining of fluorescently labeled EVs along with vesicular compartments localized predominantly in perinuclear regions 4 h after *i.n*. administration. Therefore, the nasal route is receiving considerable attention for administering drugs that net systemically. Nevertheless, it should be noted that there are several limitations including limited volume that can be dropped or sprayed into the nasal cavity, as well as removal of the drug by mucociliary clearance. Interesting, some investigators hypothesized that PD actually has its origin in the bulbs olfactory, and then spreads thought the brain ascending cell-by-cell through brainstem, midbrain, and other regions of the brain [60]. Thus, we reasoned that *i.n*. administration of EV-GDNF may work in the same manner as the natural disease spread, delivering therapeutic GDNF to the most affected brain areas.

Parkin Q311(X)A mice were treated at month four, weekly three times, and their locomotor functions were assessed over a year. Notably, the *i.n*. treatments with EV-GDNF significantly improved mobility of PD animals compared to PD groups treated with saline, or sham EVs up to the levels in healthy WT mice. Along with the improved mobility, OFA studies showed substantial improvements in behavior patterns, including decreases in the hyperactivity and anxiety-like behavior. At the endpoint, mice were sacrificed and neuroprotection effects and decreases in oxidative stress in the brain of PD animals treated with EV-GDNF were confirmed by histological evaluations. Specifically, EV-GDNF treatments produced significant neuroprotection, and reduced neuroinflammation in Parkin Q311(X)A mice. These findings could be of great importance for clinical applications of EV-based treatments. Regarding the possible mechanisms of action EV-based formulations in PD mice, GDNF is known to interact with a receptor of GDNF family, GFRA2 [61]. We hypothesized that EVs nanocarriers would protect GDNF against degradation and facilitate the delivery of this therapeutic protein to the brain tissues. Some portion of GDNF molecules may be incorporated into EVs lumen, and some of them – associated with EVs outside membranes. However, in order to produce its therapeutic effect,

GDNF should be released from EVs upon arrival to the target tissues and interact with GFRA2 receptor. Therefore, the absence of strong attachment to the EVs membrane that allows GDNF molecules release in the brain may be preferable. Of-note, superior therapeutic effects of EV-GDNF in PC12 neurons that are known to express GDNF receptor compared to high concentration of GDNF alone were shown earlier *in vitro* studies by confocal microscopy [40]. Specifically, we reported a pronounced outgrowth of axons and dendrites in neurons cultured with EV-GDNF that were greater than those caused by a high dose of commercially available GDNF.

Another crucial point is using anti-inflammatory M2-subtype of parent macrophages to further boost the therapeutic effect of EV-based formulations and ensure absence of side effects, including pro-inflammatory factors, cytokines and free radicals. We reported earlier [19] that EVs can reflect properties of their parent cells and carry the same signal molecules. For example, we demonstrated that EVs released from M2 anti-inflammatory macrophages express M2 subtype markers, Arg1 and CD206. In contrast, EVs released by M1 macrophages carry pro-inflammatory markers, such as iNOS. Of note, genetic modification of M2-macrophages did not change their subtype [19] that was confirmed on brain slides of PD mice injected with pre-transfected cells. We also report here no systemic toxicity followed EV-GDNF treatments.

Notably, EV-GDNF interventions resulted in a prolonged abrogation of neurodegeneration and neuroinflammation by EV-GDNF treatments in PD animals. We speculated that these sustained therapeutic effects may be attributed to the fact that along with GDNF, EVs released by GDNF-transfected macrophages carry genetic material encoding this therapeutic protein, GDNF-DNA. This may result in transfection of brain tissues and overexpression of GDNF that would explain these prolonged therapeutic effects. Indeed, one of natural functions of EVs is the transfer of genetic information to nearby and distant organs and tissues, including the transfer of both coding and non-coding RNAs to recipient cells [62]. We reported earlier that EVs released by pre-transfected parent cells contain the encoded therapeutic or reporter protein (*i.e*. catalase, or TPP1, or luciferase, or green fluorescence protein), along with genetic material, *p*DNA and mRNA encoding these proteins. Moreover, we found in EVs a transcription factor that was involved in the encoded gene expression (*i.e*. NF-kb) [53]. Thus, endogeniosly loaded EV-GDNF may represent a potent, non-immunogenic, naturally manufactured kit for transfection of disease tissues. We hypothesized that the accumulation of EV-GDNF in brain tissues may result in the transfer of EVs cargo to the tissues and *de novo* synthesis of the encoded protein in target cells. As such, EVs obtained from genetically modified parent cells represent a novel class of vectors that may accomplish gene transfer at the distant organs, similar to their natural functions. The mechanistic studies regarding this mechanism are ongoing in our lab and will be reported in the following publications.

Collectively our data indicates that EV-based formulations of GDNF is a promising therapeutic modality that can provide a versatile and potent strategy for CNS delivery of therapeutics, which can be applied to different neurodegenerative disorders. Overall, the exploration of mechanisms involved in targeted CNS transport of EV-based drug formulations is crucial for their therapeutic applications and transformation the field of precision medicine.

## Supporting information

All Supplementary Figures and Tables

## Supplementary Materials

The following are available online at www.mdpi.com/xxx/s1, Figure S1: Absence of gross toxicity of EV-GDNF treatment in Parkin Q311(X)A mice, Table S1: Operation parameters of parent macrophages electroporation upon transfection with GDNF-encoding *p*DNA, Table 2: Primary antibodies used for Simple Western Blot, Table S3: Expression levels of EV-specific proteins in EVs released by GDNF-transfected macrophages, Table S4: Murine integrin proteotypic tryptic peptides detectable for the label free quantitative assessment, Table S5: Effect of EV-GDNF on inflammation and neurodegeneration in PD mice.

## Author Contributions

Conceptualization, E.V.B.; methodology, E.V.B., P.C.S., E.B.H., and N.E-H.; validation, Y.Z., M.J.H., J.K.F., Y.S.J., C.J.A., C.J.S., and M.R.; data curation, M.J.H., and E.B.H.; writing—original draft preparation, E.V.B, E.B.H. and N.E-H.; writing—review and editing, E.V.B. and M.S.L. All authors have read and agreed to the published version of the manuscript.

## Institutional Review Board Statement

The study was conducted according to the guidelines of the Declaration of Helsinki and approved by the Institutional Animal Care and Use Committee of University of North Carolina at Chapel Hill (protocol code 15-141.0-B, approval date 04/2017).

## Acknowledgments

We are very grateful to Mr. and Mrs. Lehrman and Mr. T. Greenwood for financial support and various invaluable comments and suggestions. We would like to acknowledge the support of the UNC Nanomedicines Characterization Core Facility (http://ncore.web.unc.edu) in the EVs characterization. We are also thankful to the Senior Director of Development at UNC Kelly Collins for her assistance with communication the strategy and facilitating the support of this project.

## Appendix A

**Supplementary Figure S1.**
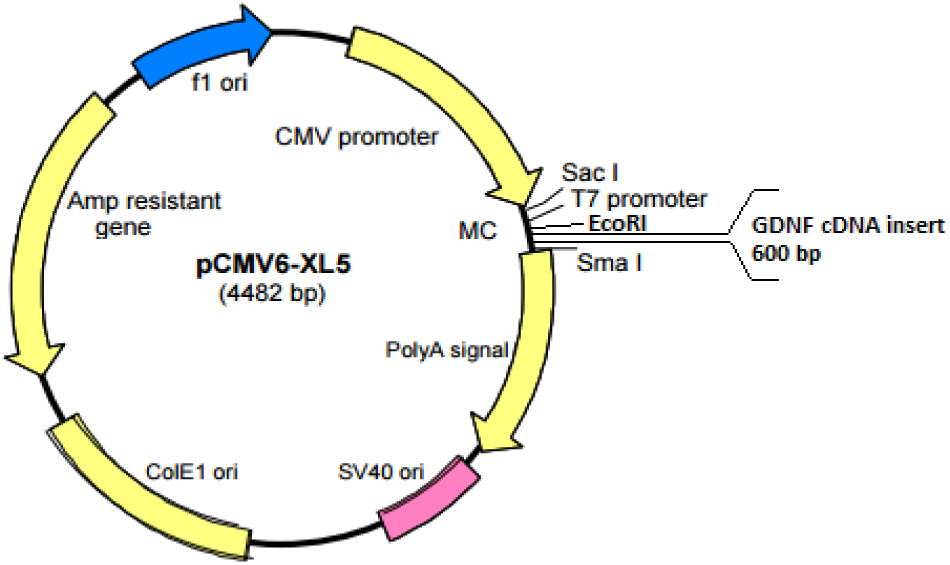
Plasmid map for GDNF production.

**Supplementary Figure S2.**
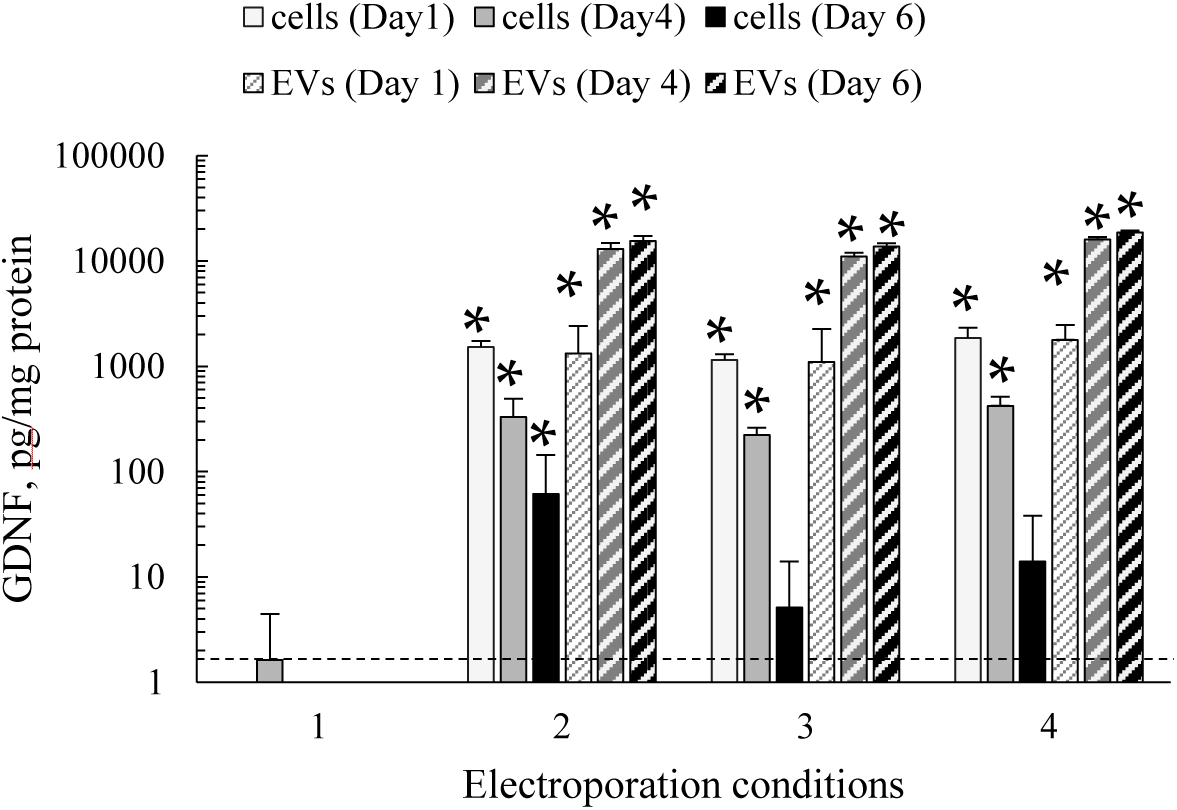
Transfection of primary murine macrophages with GDNF-encoding *p*DNA. Bone-marrow derived macrophages were transfected by electroporation using four different conditions described in Experimental section. Then, cells were washed and cultured in complete media for up to 6 days. The GDNF expression levels in cells (solid bars), and EVs collected from conditioned media (stripped bars) was assessed by ELISA on day 1 (white bars), day 4 (grey bars), and day 6 (black bars). Successful transfection was accomplished with three conditions (#2 - #4). N = 4, *p < 0.05, compared to sham-transfected macrophages (dashed line, condition #1).

**Supplementary Figure S3.**
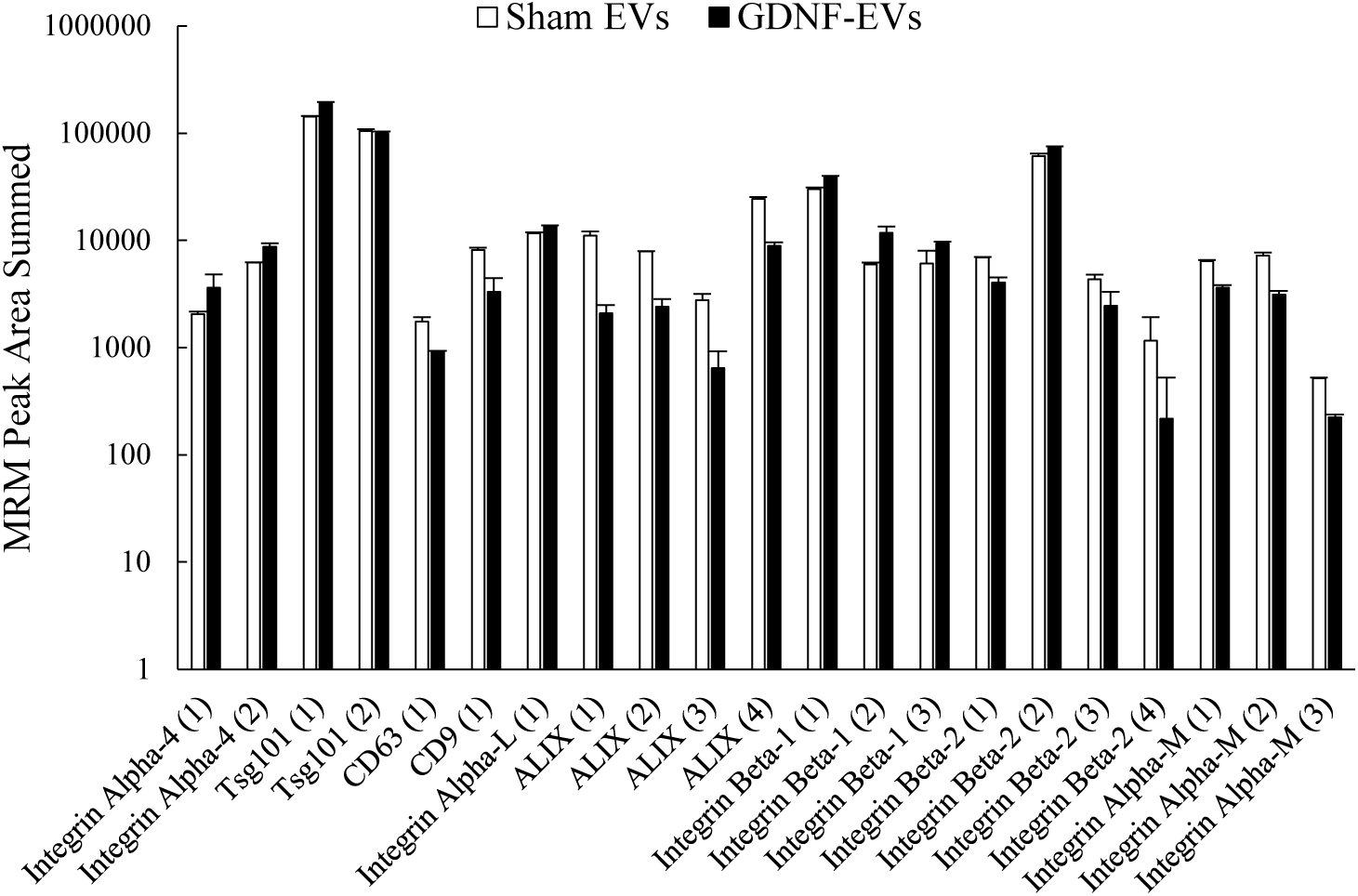
Quantification of Integrins and Tetraspanins in EVs by Label Free Targeted Quantitative Proteomics. EVs samples from sham-transfected (white bars), and GDNF-transfected (black bars) macrophages were digested (*N* =3) with trypsin and examined by nano-liquid chromatography tandem MS (nanoLC–MS/MS) with multiple reaction monitoring (MRM). Samples of 20 µg total protein were used, and 0.08 µg (0.4 % of the sample) was injected. No significant differences in specific proteins expression were found between sham EVs and EV-GDNF (t-tests, *p* < 0.05). Peptide identification is shown in **Supplementary Table S3**. A CD81 peptide employed in other studies was not detected in these analyses. Values are means ± SD.

**Supplementary Figure S4.**
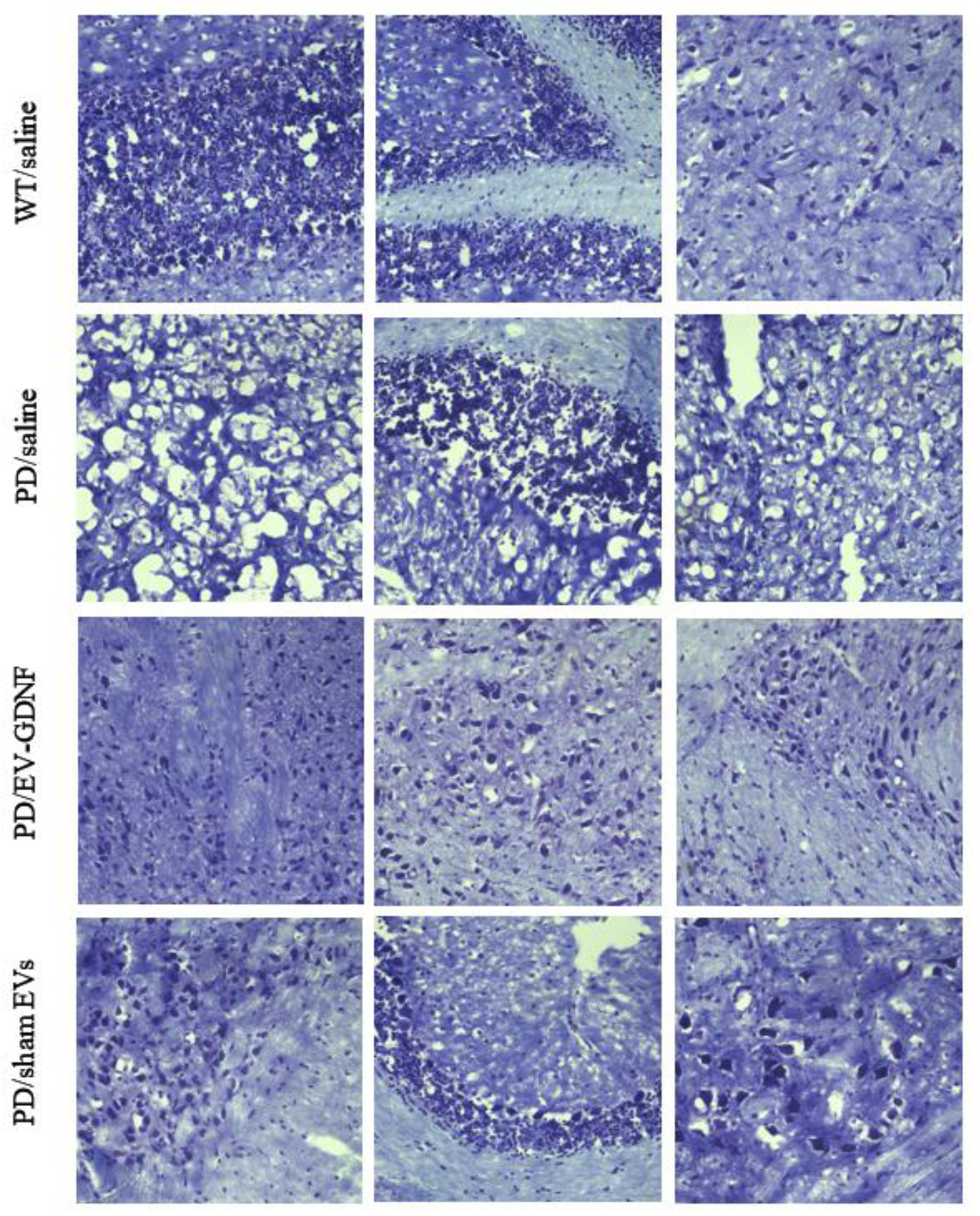
Histological analysis of neuroprotective effects by EV-GDNF in Parkin Q311(X)A mice. Transgenic mice (4 mo. old) were intranasally injected with: saline (10 µL/mouse), or EV-GDNF (3×10^9^ particles/10 µL/mouse), or sham EVs (3×10^9^ particles/10 µL/mouse) weekly three times. Wild type control mice were intranasally injected with saline (10 µL/mouse). Animals were sacrificed at mo. 16, brain slides were stained with Nissl staining. The obtained bright light images show lower number of Nissl bodies with neuronal shrinkage and damages tissues with degeneration in the neurons in PD mice treated with saline when compared to WT mice. Histological analysis indicates neuroprotective effects in the brain of PD mice treated with GDNF-EVs with healthy morphology in tissue structure and high integrity of neurons when comparted to PD mice treated with saline. The administration of sham EVs did not have significant therapeutic effect in PD mice.

**Supplementary Figure S5.**
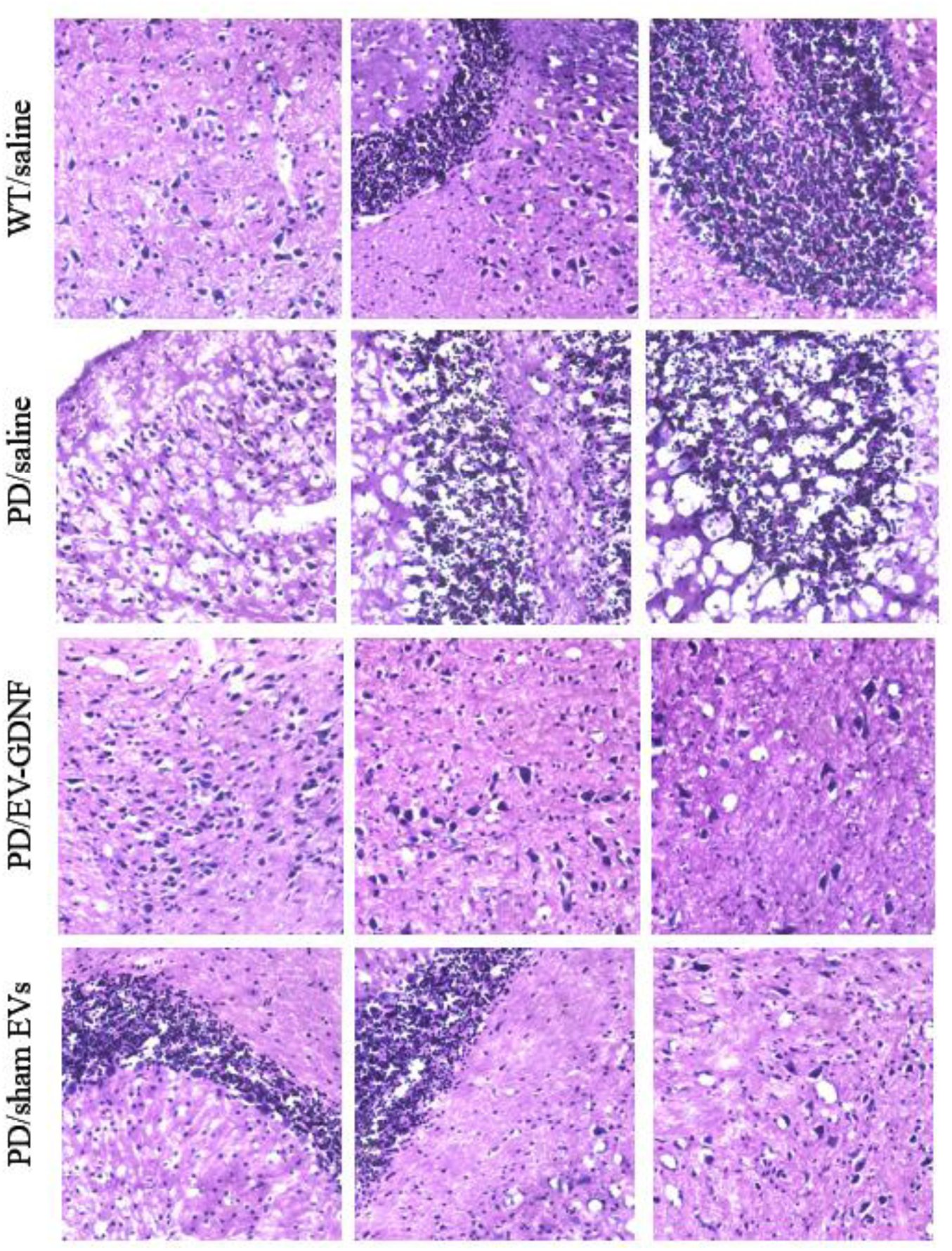
Histological analysis of neuroprotective effects by EV-GDNF in Parkin Q311(X)A mice. Transgenic mice (4 mo. old) were intranasally injected with: saline (10 µL/mouse), or EV-GDNF (3×10^9^ particles/10 µL/mouse), or sham EVs (3×10^9^ particles/10 µL/mouse) weekly three times. Wild type control mice were intranasally injected with saline (10 µL/mouse). Animals were sacrificed at mo. 16, brain slides were stained with Nissl staining. The obtained bright light images show damaged tissues with degeneration in the neurons in PD mice treated with saline when compared to WT mice. Histological analysis indicates neuroprotective effects in the brain of PD mice treated with GDNF-EVs with healthy morphology in tissue structure when comparted to PD mice treated with saline. The administration of sham EVs did not have significant therapeutic effect in PD mice.

**Supplementary Figure S6.**
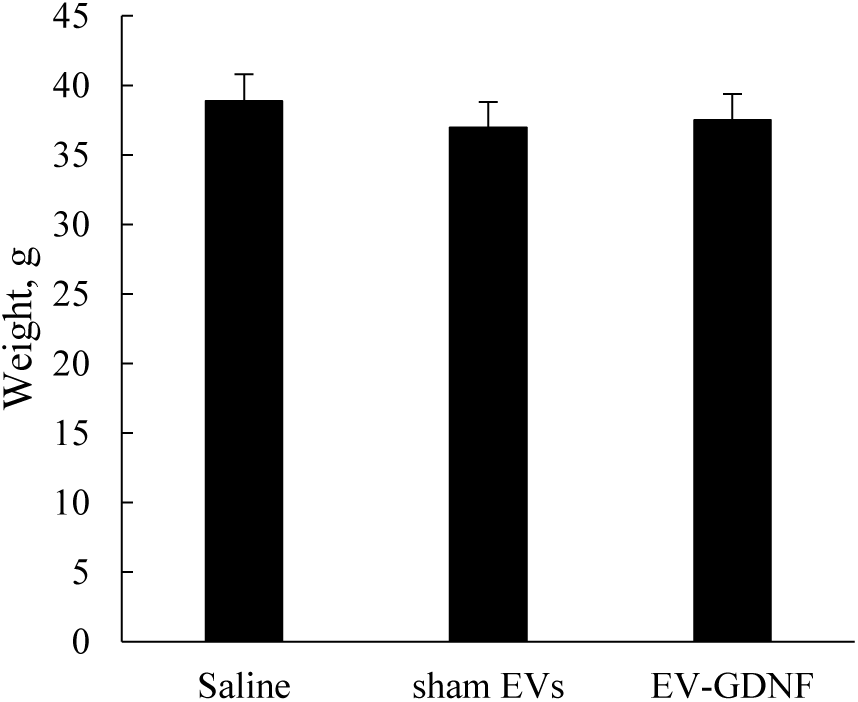
Absence of gross toxicity of EV-GDNF treatment in Parkin Q311(X)A mice. Transgenic mice (4 mo. of age) were *i.n*. injected with saline, or EV-GDNF, or sham EVs (3×10^9^ particles/10 µL/mouse, once a week, 3x weeks). At 16 mo. of age total weigh of the animals was recorded. No gross toxicity manifested in the losing weight was detected in mice injected with EV-GDNF and well as sham EVs.

**Supplementary Table S1.**
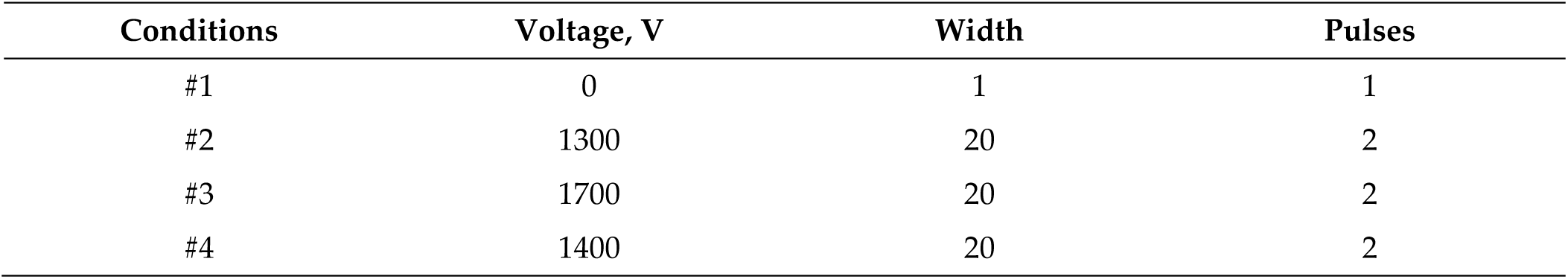
Operation parameters of parent macrophages electroporation upon transfection with GDNF-encoding *p*DNA.

**Supplementary Table S2.**
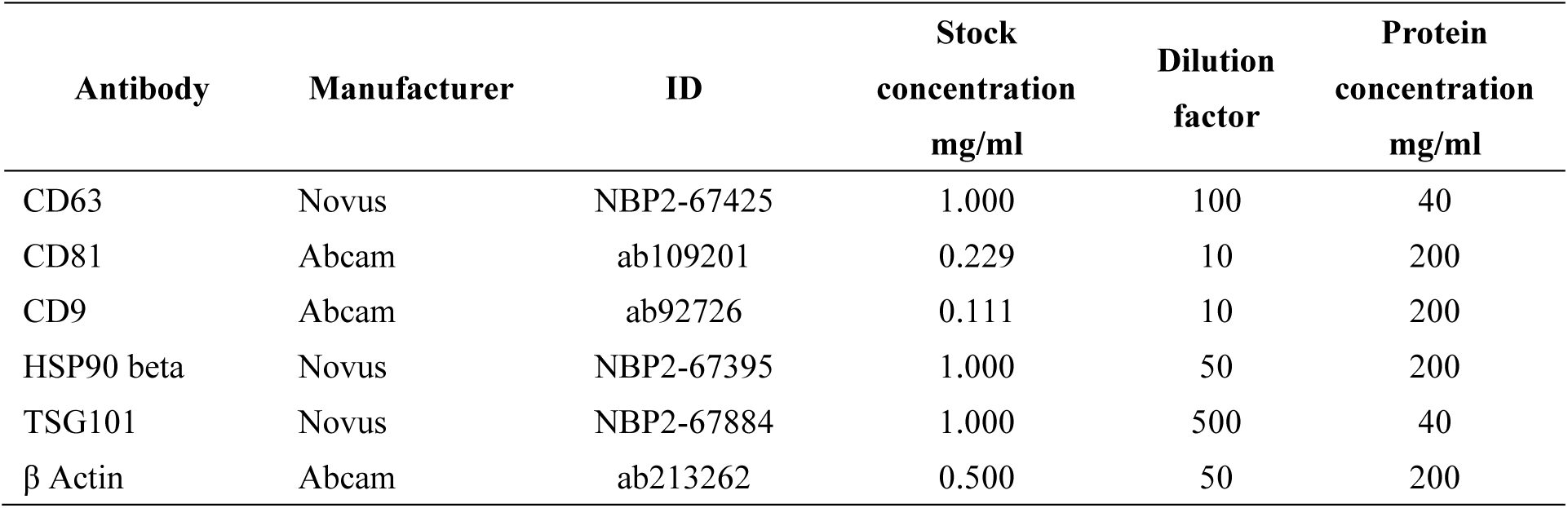
Primary antibodies used for Simple Western Blot.

**Supplementary Table S3.**
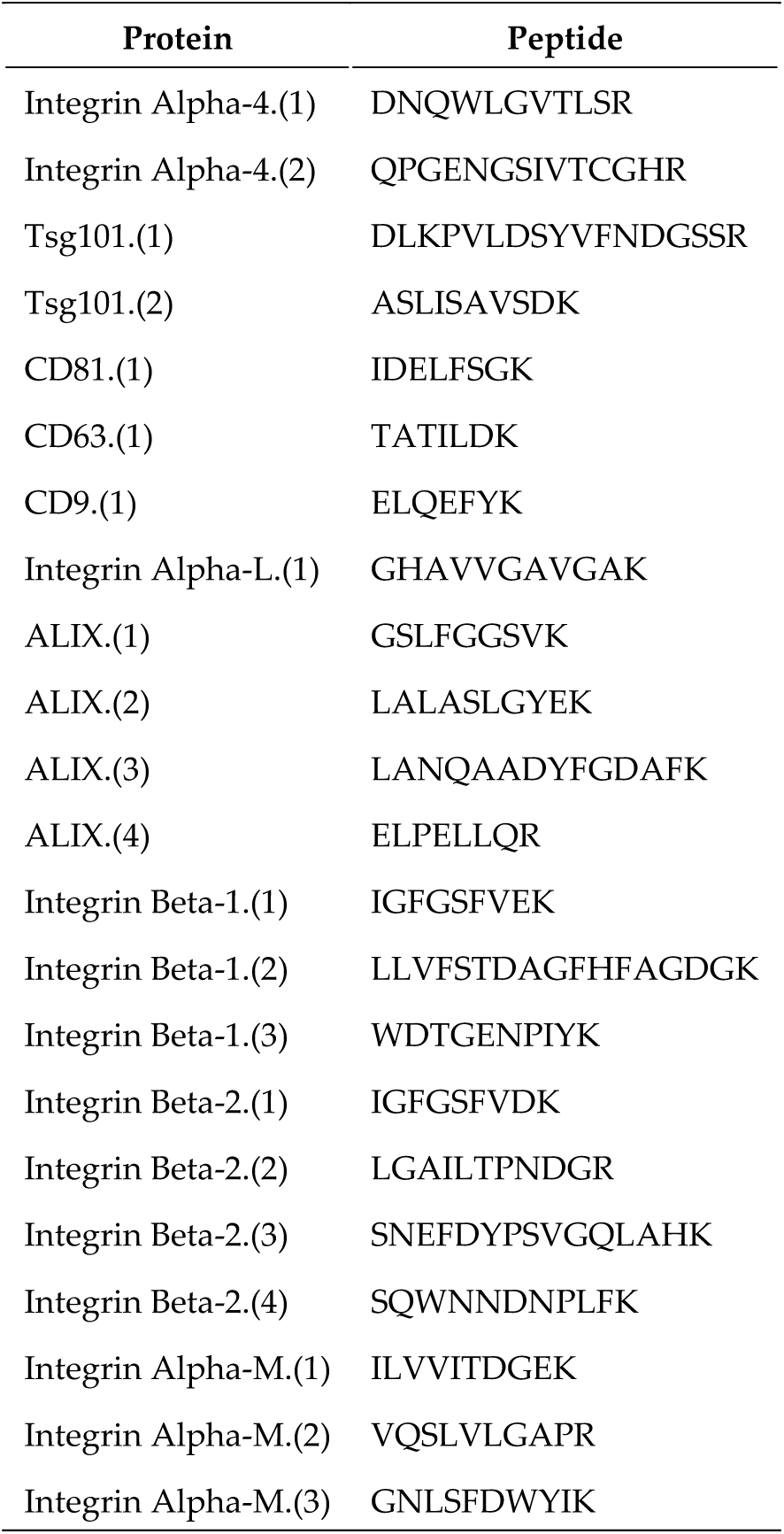
Murine integrin proteotypic tryptic peptides detectable for the label free quantitative assessment.

**Supplementary Table S4.**
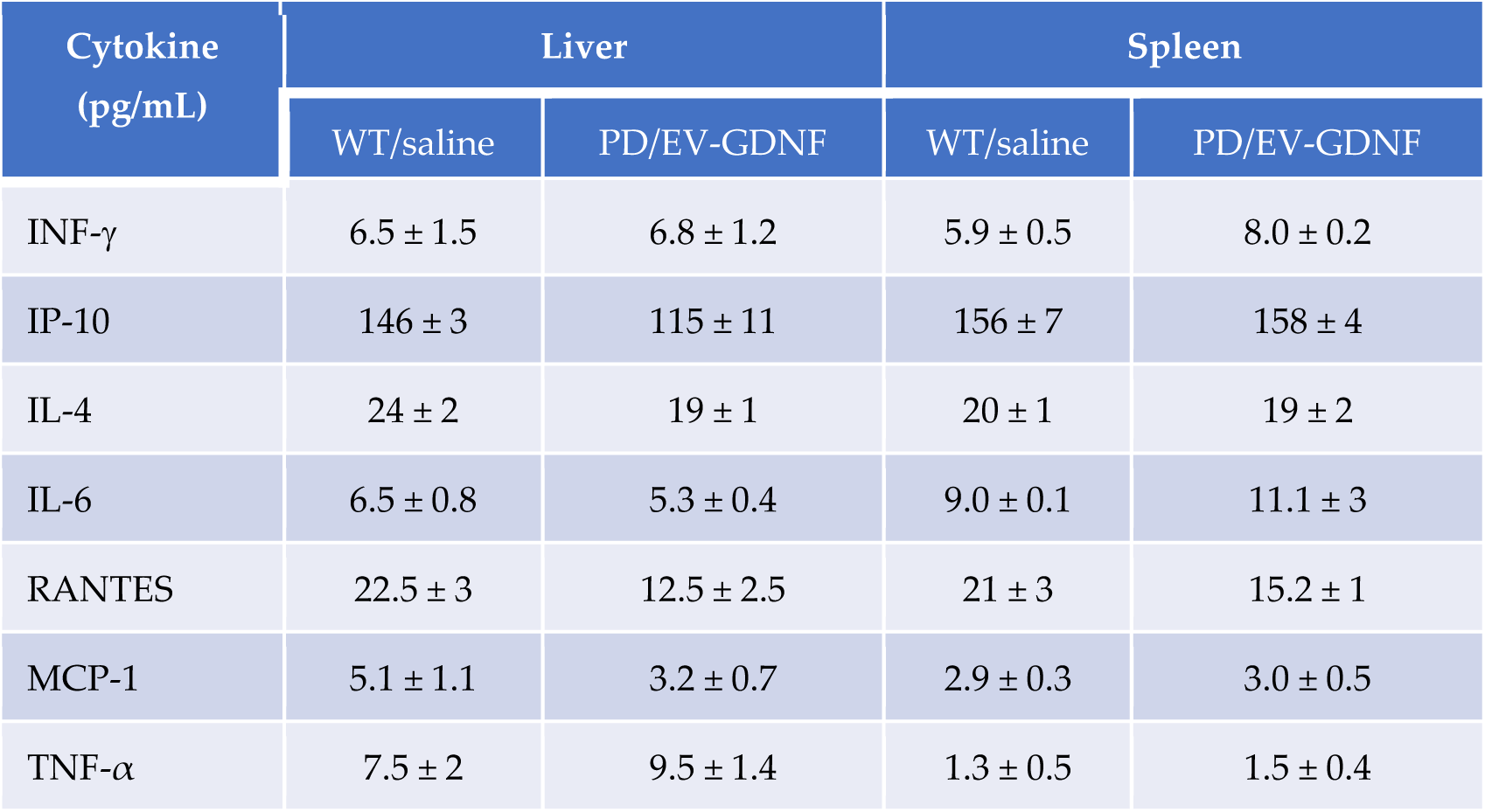
Effect of EV-GDNF on the expression of pro-inflammatory cytokines.

| Cytokine<br>(pg/mL) | Liver |  | Spleen |  |
| --- | --- | --- | --- | --- |
|  | WT/saline | PD/EV-GDNF | WT/saline | PD/EV-GDNF |
| INF-γ | 6.5 ± 1.5 | 6.8 ± 1.2 | 5.9 ± 0.5 | 8.0 ± 0.2 |
| IP-10 | 146 ± 3 | 115 ± 11 | 156 ± 7 | 158 ± 4 |
| IL-4 | 24 ± 2 | 19 ± 1 | 20 ± 1 | 19 ± 2 |
| IL-6 | 6.5 ± 0.8 | 5.3 ± 0.4 | 9.0 ± 0.1 | 11.1 ± 3 |
| RANTES | 22.5 ± 3 | 12.5 ± 2.5 | 21 ± 3 | 15.2 ± 1 |
| MCP-1 | 5.1 ± 1.1 | 3.2 ± 0.7 | 2.9 ± 0.3 | 3.0 ± 0.5 |
| TNF-α | 7.5 ± 2 | 9.5 ± 1.4 | 1.3 ± 0.5 | 1.5 ± 0.4 |

