## Supplementary material for "Using Extracellular Vesicles Released by GDNF-transfected Macrophages for Therapy of Parkinson’s Disease": All Supplementary Figures and Tables

### Supplementary Information

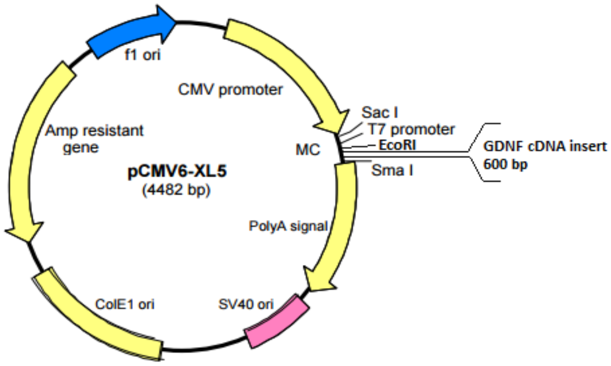


**Supplementary Figure S1. Plasmid map for GDNF production**

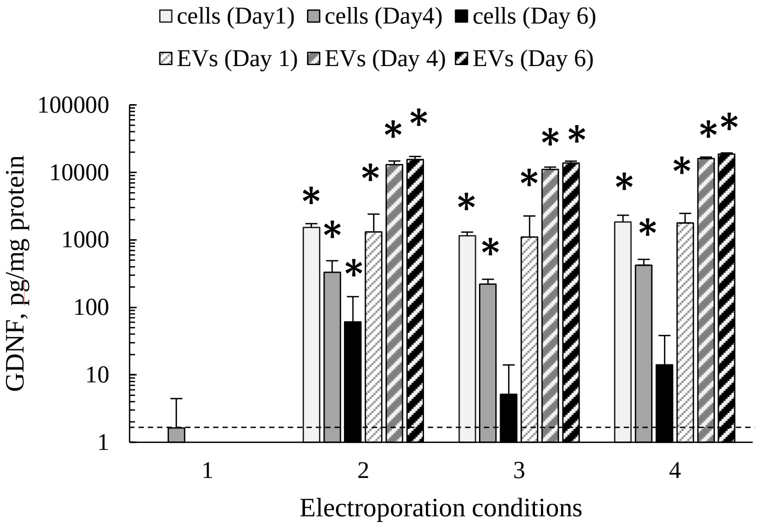


**Supplementary Figure S2. Transfection of primary murine macrophages with GDNF-encoding *p*DNA.** Bone-marrow derived macrophages were transfected by electroporation using four different conditions described in Experimental section. Then, cells were washed and cultured in complete media for up to 6 days. The GDNF expression levels in cells (solid bars), and EVs collected from conditioned media (stripped bars) was assessed by ELISA on day 1 (white bars), day 4 (grey bars), and day 6 (black bars). Successful transfection was accomplished with three conditions (#2 - #4). N = 4, *p < 0.05, compared to sham-transfected macrophages (dashed line, condition #1).

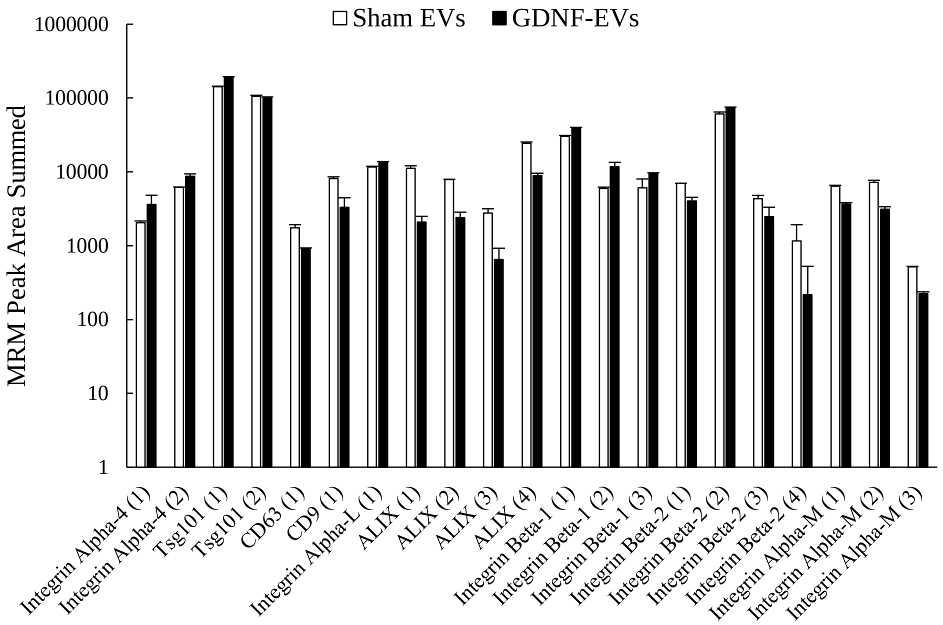


**Supplementary Figure S3. Quantification of Integrins and Tetraspanins in EVs by Label Free Targeted Quantitative Proteomics.** EVs samples from sham-transfected (white bars), and GDNF-transfected (black bars) macrophages were digested ($N$ =3) with trypsin and examined by nano-liquid chromatography tandem MS (nanoLC–MS/MS) with multiple reaction monitoring (MRM). Samples of 20 µg total protein were used, and 0.08 µg (0.4 % of the sample) was injected. No significant differences in specific proteins expression were found between sham EVs and EV-GDNF (t-tests, $p$ < 0.05). Peptide identification is shown in **Supplementary Table S3**. A CD81 peptide employed in other studies was not detected in these analyses. Values are means ± SD.

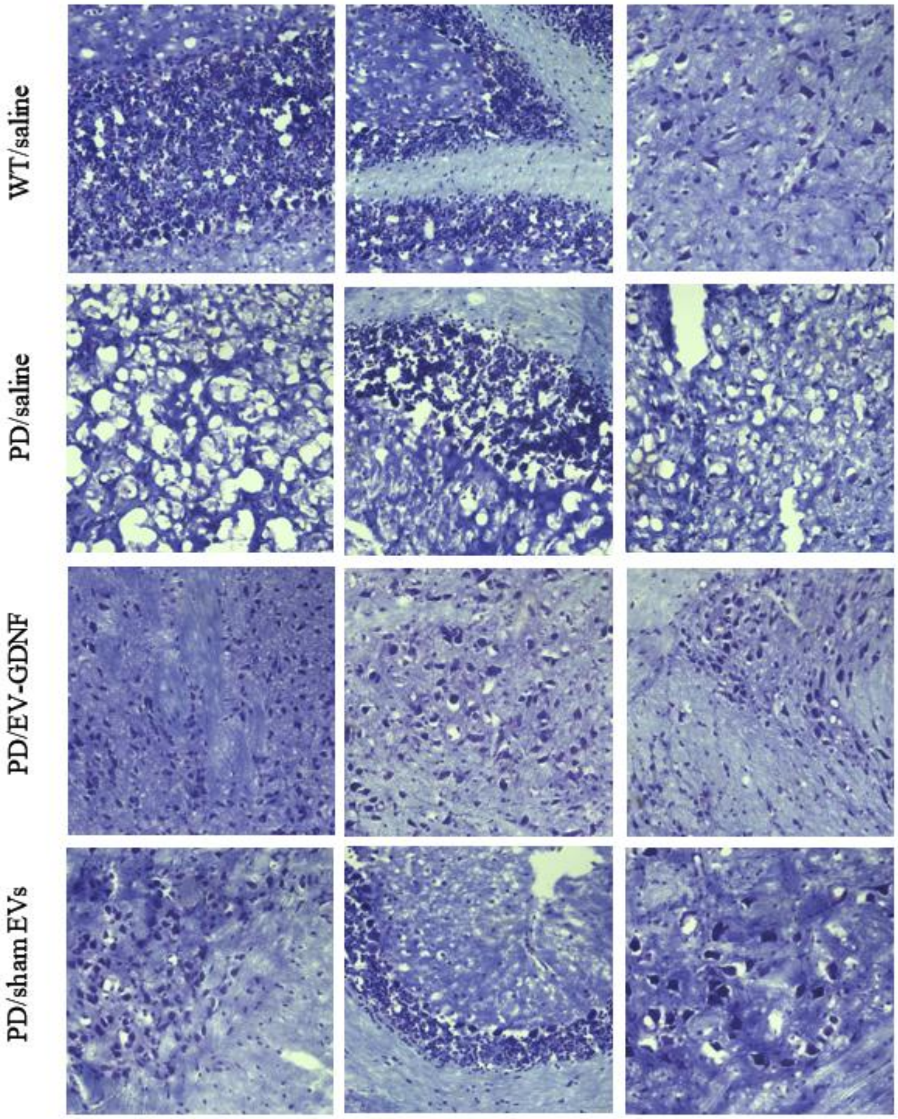


**Supplementary Figure S4. Histological analysis of neuroprotective effects by EV-GDNF in Parkin Q311(X)A mice.** Transgenic mice (4 mo. old) were intranasally injected with: saline (10 µL/mouse), or EV-GDNF ($3x10^9$ particles/10 µL/mouse), or sham EVs ($3x10^9$ particles/10 µL/mouse) weekly three times. Wild type control mice were intranasally injected with saline (10 µL/mouse). Animals were sacrificed at mo. 16, brain slides were stained with Nissl staining. The obtained bright light images show lower number of Nissl bodies with neuronal shrinkage and damages tissues with degeneration in the neurons in PD mice treated with saline when compared to WT mice. Histological analysis indicates neuroprotective effects in the brain of PD mice treated with GDNF-EVs with healthy morphology in tissue structure and high integrity of neurons when comparted to PD mice treated with saline. The administration of sham EVs did not have significant therapeutic effect in PD mice.

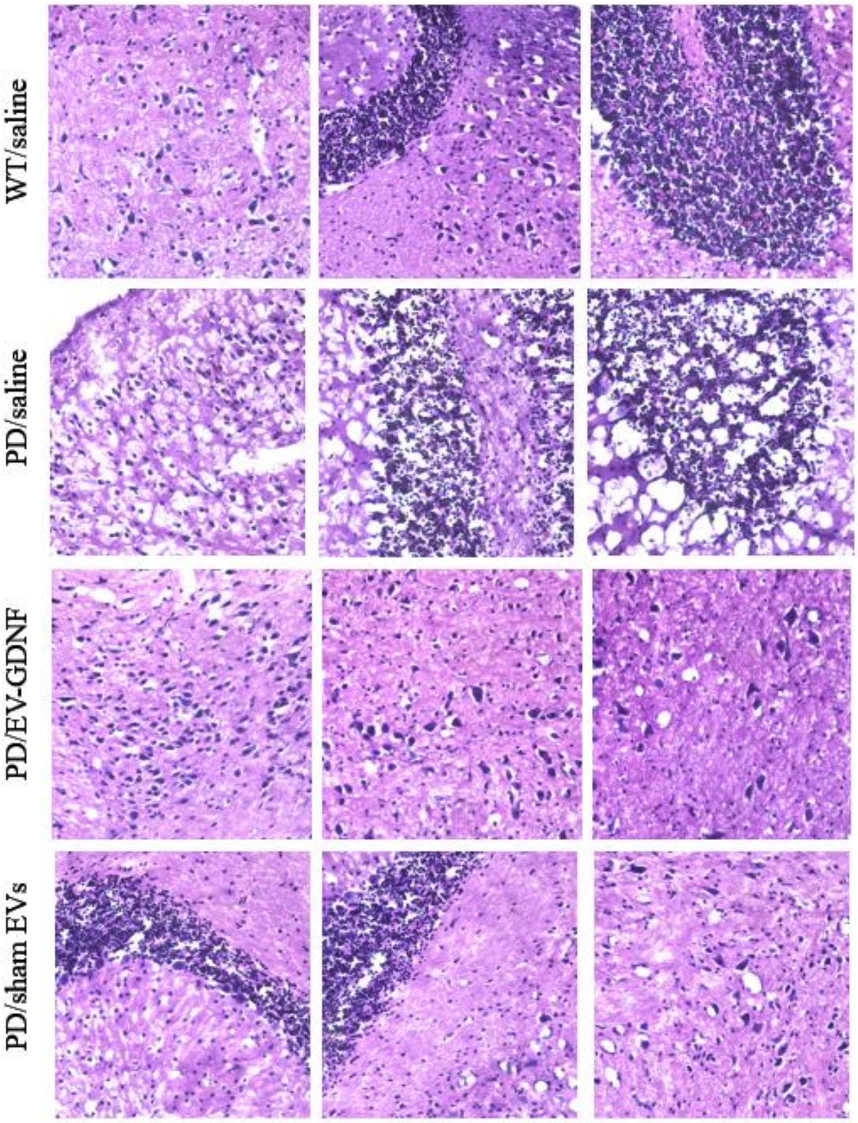


**Supplementary Figure S5. Histological analysis of neuroprotective effects by EV-GDNF in Parkin Q311(X)A mice.** Transgenic mice (4 mo. old) were intranasally injected with: saline (10 µL/mouse), or EV-GDNF ($3x10^9$ particles/10 µL/mouse), or sham EVs ($3x10^9$ particles/10 µL/mouse) weekly three times. Wild type control mice were intranasally injected with saline (10 µL/mouse). Animals were sacrificed at mo. 16, brain slides were stained with Nissl staining. The obtained bright light images show damaged tissues with degeneration in the neurons in PD mice treated with saline when compared to WT mice. Histological analysis indicates neuroprotective effects in the brain of PD mice treated with GDNF-EVs with healthy morphology in tissue structure when comparted to PD mice treated with saline. The administration of sham EVs did not have significant therapeutic effect in PD mice.

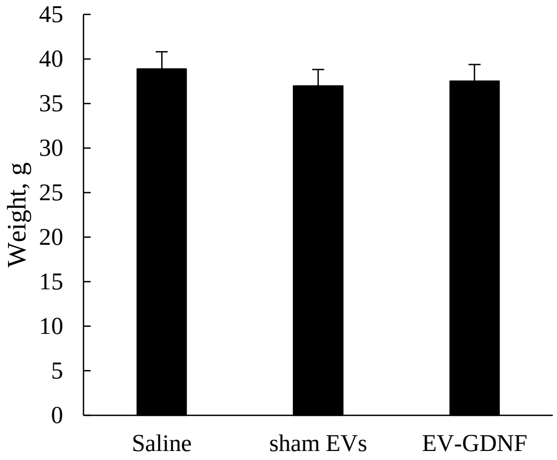


**Supplementary Figure S6. Absence of gross toxicity of EV-GDNF treatment in Parkin Q311(X)A mice**. Transgenic mice (4 mo. of age) were *i.n.* injected with saline, or EV-GDNF, or sham EVs ($3x10^9$ particles/10 µL/mouse, once a week, 3x weeks). At 16 mo. of age total weigh of the animals was recorded. No gross toxicity manifested in the losing weight was detected in mice injected with EV-GDNF and well as sham EVs.

**Supplementary Table S1. Operation parameters of parent macrophages electroporation upon transfection with GDNF-encoding *p*DNA**

| Conditions | Voltage, V | Width | Pulses |
|---|---|---|---|
| #1 | 0 | 1 | 1 |
| #2 | 1300 | 20 | 2 |
| #3 | 1700 | 20 | 2 |
| #4 | 1400 | 20 | 2 |

**Supplementary Table S2. Primary antibodies used for Simple Western Blot**

| Antibody | Manufacturer | ID | Stock concentration mg/ml | Dilution factor | Protein concentration mg/ml |
|---|---|---|---|---|---|
| CD63 | Novus | NBP2-67425 | 1.000 | 100 | 40 |
| CD81 | Abcam | ab109201 | 0.229 | 10 | 200 |
| CD9 | Abcam | ab92726 | 0.111 | 10 | 200 |
| HSP90 beta | Novus | NBP2-67395 | 1.000 | 50 | 200 |
| TSG101 | Novus | NBP2-67884 | 1.000 | 500 | 40 |
| β Actin | Abcam | ab213262 | 0.500 | 50 | 200 |

**Supplementary Table S3. Murine integrin proteotypic tryptic peptides detectable for the label free quantitative assessment.**

| Protein | Peptide |
|---|---|
| Integrin Alpha-4.(1) | DNQWLGVTLSR |
| Integrin Alpha-4.(2) | QPGENGSIVTCGHR |
| Tsg101.(1) | DLKPVLDSYVFNDGSSR |
| Tsg101.(2) | ASLISAVSDK |
| CD81.(1) | IDELFSGK |
| CD63.(1) | TATILDK |
| CD9.(1) | ELQEFYK |
| Integrin Alpha-L.(1) | GHAVVGAVGAK |
| ALIX.(1) | GSLFGGSVK |
| ALIX.(2) | LALASLGYEK |
| ALIX.(3) | LANQAADYFGDAFK |
| ALIX.(4) | ELPELLQR |
| Integrin Beta-1.(1) | IGFGSFVEK |
| Integrin Beta-1.(2) | LLVFSTDAGFHFAGDGK |
| Integrin Beta-1.(3) | WDTGENPIYK |
| Integrin Beta-2.(1) | IGFGSFVDK |
| Integrin Beta-2.(2) | LGAILTPNDGR |
| Integrin Beta-2.(3) | SNEFDYPSVGQLAHK |
| Integrin Beta-2.(4) | SQWNNDNPLFK |
| Integrin Alpha-M.(1) | ILVVITDGEK |
| Integrin Alpha-M.(2) | VQSLVLGAPR |
| Integrin Alpha-M.(3) | GNLSFDWYIK |

**Supplementary Table S4. Effect of EV-GDNF on the expression of pro-inflammatory cytokines**

| Cytokine (pg/mL) | Liver | | Spleen | |
|---|---|---|---|---|
| | WT/saline | PD/EV-GDNF | WT/saline | PD/EV-GDNF |
| INF-γ | 6.5 ± 1.5 | 6.8 ± 1.2 | 5.9 ± 0.5 | 8.0 ± 0.2 |
| IP-10 | 146 ± 3 | 115 ± 11 | 156 ± 7 | 158 ± 4 |
| IL-4 | 24 ± 2 | 19 ± 1 | 20 ± 1 | 19 ± 2 |
| IL-6 | 6.5 ± 0.8 | 5.3 ± 0.4 | 9.0 ± 0.1 | 11.1 ± 3 |
| RANTES | 22.5 ± 3 | 12.5 ± 2.5 | 21 ± 3 | 15.2 ± 1 |
| MCP-1 | 5.1 ± 1.1 | 3.2 ± 0.7 | 2.9 ± 0.3 | 3.0 ± 0.5 |
| TNF-α | 7.5 ± 2 | 9.5 ± 1.4 | 1.3 ± 0.5 | 1.5 ± 0.4 |
